# Nationwide genome surveillance of carbapenem-resistant *Pseudomonas aeruginosa* in Japan

**DOI:** 10.1101/2023.12.17.572044

**Authors:** Hirokazu Yano, Wataru Hayashi, Sayoko Kawakami, Sadao Aoki, Eiko Anzai, Hui Zuo, Norikazu Kitamura, Aki Hirabayashi, Toshiki Kajihara, Shizuo Kayama, Yo Sugawara, Koji Yahara, Motoyuki Sugai

## Abstract

Japan is a country with an approximate 10 % prevalence rate of carbapenem-resistant *Pseudomonas aeruginosa* (CRPA). Currently, a comprehensive overview of the genotype and phenotype patterns of CRPA in Japan is lacking. Herein, we conducted genome sequencing and quantitative antimicrobial susceptibility testing for 382 meropenem-resistant CRPA isolates that were collected from 78 hospitals across Japan from 2019 to 2020. CRPA exhibited susceptibility rates of 52.9%, 26.4%, and 88.0% against piperacillin-tazobactam, ciprofloxacin, and amikacin, respectively, whereas 27.7% of CRPA isolates were classified as difficult-to-treat resistance *P. aeruginosa*. Of the 148 sequence types detected, ST274 (9.7%) was predominant, followed by ST235 (7.6%). The proportion of urine isolates in ST235 was higher than that in other STs (*P* = 0.0056, chi-square test). Only 4.1% of CRPA isolates carried the carbapenemase genes: *bla*_GES_ (2) and *bla*_IMP_ (13). One ST235 isolate carried the novel *bla*_IMP_ variant *bla*_IMP-98_ in the chromosome. Regarding chromosomal mutations, 87.1% of CRPA isolates possessed inactivating or other resistance mutations in *oprD*, and 28.8% showed mutations in the regulatory genes (*mexR, nalC*, and *nalD*) for the MexAB-OprM efflux pump. Additionally, 4.7% of CRPA isolates carried a resistance mutation in the PBP3-encoding gene *ftsI*. The findings from this study and other surveillance studies collectively demonstrate that CRPA exhibits marked genetic diversity and that its multidrug resistance in Japan is less prevailed than in other regions. This study contributes a valuable dataset that addresses a gap in genotype/phenotype information regarding CRPA in the Asia–Pacific region, where the epidemiological background markedly differs between regions.

## INTRODUCTION

*Pseudomonas aeruginosa* is a common opportunistic pathogen associated with pneumonia, urinary tract infections, and bacteremia in clinical settings worldwide. Currently, it ranks among the top five bacterial pathogens associated with human death globally (1). It also intrinsically shows reduced susceptibility to many antibiotics due to its possession of four types of RND family efflux pump, chromosomally encoded AmpC β-lactamase (also called *Pseudomonas*-derived class C β-lactamase: PDC), and other antimicrobial-inactivating enzymes (2, 3). Furthermore, *P. aeruginosa* can acquire resistance mutations at multiple genomic locations and antimicrobial resistance genes via mobile genetic elements, such as plasmids and integrative and conjugative elements (ICEs), after which it elicits high levels of resistance to multiple antimicrobials, thereby limiting treatment options for infected patients (3–5).

Carbapenems represent a crucial class of therapeutic options for treating *P. aeruginosa* infections. In 2017, the World Health Organization (WHO) listed carbapenem-resistant *P. aeruginosa* (CRPA) as one of the bacterial priority pathogens classified in “Priority 1: CRITICAL,” for which new effective antibiotics are urgently required (6). Horizontally transferrable carbapenemases represent a significant public health concern due to their insensitivity to the conventionally used β-lactamase inhibitors (7). A recent cohort study demonstrated a 9% higher 30-day mortality rate among patients infected with carbapenemase-producing CRPA when compared to those infected with carbapenemase-nonproducing CRPA (8). Currently, three classes of β-lactamases, namely class A, class B (known as metallo-β-lactamases or MBLs), and class D β-lactamases, are recognized as transferrable carbapenemase. Globally disseminated carbapenemase types in *P. aeruginosa* include VIM, IMP, NDM, GES, and KPC (9). In addition to the horizontal acquisition of carbapenemases, *P. aeruginosa* can develop carbapenem resistance through the mutational inactivation of several chromosomally coded functions. These include the inactivation of the OprD porin (10) and the inactivation of two direct or one indirect transcriptional regulators (repressors, MexR, NalC, and NalD) of the MexAB-OprM effux pump (11–13). Furthermore, an experimental evolution study revealed that nonsynonymous substitutions in *ftsI*, an essential gene encoding PBP3, contribute to carbapenem resistance (14).

Recently, international consortia have made efforts to build a large *P. aeruginosa* dataset to obtain quantitative information for CRPA prevalence, its susceptibility profiles, and the possession rate of carbapenemases (8, 15–17); Notably, in the Asia–Pacific region, the Antimicrobial Testing Leadership and Surveillance (ATLAS) program took the initiative of *P. aeruginosa* surveillance, revealing distinct patterns of resistance rate and predominant β-lactamase levels in the *P. aeruginosa* population between 12 countries/regions, but the ATLAS did not cover genome surveillance (16). Hence, information of the major types of resistance mechanisms and dominant clones of *P. aeruginosa* in the Asia–Pacific region remain lacking.

Japan exhibits characteristic features in the epidemiology of antimicrobial-resistant bacterial infection in the Asia–Pacific region. Regarding carbapenem resistance, the prevalence of meropenem-resistant *E. coli* in bloodstream infections in Japan as of 2019 was 2 per 100,000 test patients, which is lower than, for example, 79 per 100,000 test patients observed in Singapore, which is located in Southeast Asia, according to the WHO Global Antimicrobial Resistance and Use Surveillance System (18). The established national phenotypic antimicrobial resistance surveillance program—the Japan Nosocomial Infections Surveillance (JANIS) (19, 20)—reported that the carbapenem (meropenem) resistance rate of *P. aeruginosa* in hospitalized patients in Japan, as of 2020, was 10.5%. This was average among representative countries where antimicrobial resistance surveillance for *P. aeruginosa* was once or is currently being performed; the resistance rate was 29.3% in India (ATLAS 2015–2019), 19.3% in China (CHINET 2020), 6.5% in Australia (ATLAS 2015–2019), 28.5% in Poland (EARS-net 2020), 15.7% in the Czech Republic (EARS-net 2020), 13.9% in Germany (EARS-net 2020), 12.6% in France (EARS-net 2020), and 3.7% in Finland (EARS-net 2020) (16, 21, 22).

To obtain an overview of both genomic features and resistance phenotypes of bacterial pathogens in Japan, we initiated the national bacterial genome surveillance—Japan Antimicrobial-Resistant Bacterial Surveillance (JARBS)—in 2019 (23). Through the JARBS-PA project performed in 2019 and 2020, 382 meropemen-resistant CRPA isolates were collected from 78 hospitals across Japan. Herein, we report the genotype and susceptibility patterns of the CRPA isolates in Japan based on genome surveillance and standardized MIC remeasurements.

## RESULTS

### Nationwide sampling

From 2019 to 2020, more than 600 *P. aeruginosa* isolates were transferred from Japanese hospitals to the Antimicrobial Resistance Research Center (AMR-RC) at the National Institute of Infectious Diseases (NIID) based on the following criterion: meropenem or imipenem MIC ≥8μg/ml in initial testing at the isolation site. All the collected isolates were tested for antipseudomonal agent MICs using the Beckman MicroScan system at AMR-RC. In total, 382 *P. aeruginosa* isolates showing meropenem MIC ≥8μg/ml in the tests at AMR-RC were defined as CRPA and focused on in this study. The 382 CRPA isolates originated from nonoverlapping patients in 78 hospitals (the number of collected isolates per island was as follows: *n* = 30 from Hokkaido Island, *n* = 307 from Honshu Island, *n* = 7 from Shikoku Island, *n* = 32 from Kyusyu Island, and *n* = 6 from Okinawa Island). Source specimens were classified into one of the following JANIS categories (19, 20): Oral/Endotracheal/Respiratory (*n* = 189), Urinary/Genital (*n* = 76), Blood/Fluid (*n* = 40), Digestive (*n* = 27), and Other (*n* = 50). Table S1 provides more detailed source categories and metadata for each isolate. Draft genome sequences of 382 CRPA strains were determined to identify the sequence type (ST), resistance genes, and mutations.

### Antimicrobial susceptibility profiles and STs of CRPA in Japan

Table 1 summarizes the susceptibilities for antipseudomonal agents. Although the collected CRPA isolates were often nonsusceptible to antipseudomonal agents other than meropenem, the percentage of susceptible isolates remained 52.9% for piperacillin-tazobactam, 26.4% for ciprofloxacin, and 88.0% for amikacin based on breakpoints in CLSI M100-ed 32 in 2022 (Table 1). When focused on isolates from urine (CLSI M100-ed 33 in 2023), the proportion of amikacin-susceptible isolates was decreased to 75%.

**TABLE 1.** Antimicrobial susceptibilities of 382 meropenem-resistant CRPA isolates.

| Antimicrobial agents | Percentage of susceptibility group<br>(number of isolates)<sup>a</sup> |  |  |
| --- | --- | --- | --- |
|  | S | I | R |
| Antipseudomonal carbapenems |  |  |  |
| Imipenem | 1.0 (4) | 2.9 (11) | 96.1 (367) |
| Meropenem | 0.0 (0) | 0.0 (0) | 100 (382) |
| Doripenem | 10.7 (41) | 36.6 (136) | 53.7 (205) |
| Antipseudomonal cephalosporins |  |  |  |
| Ceftazidime | 53.1 (203) | 12.8 (49) | 34.0 (130) |
| Cefepime | 47.4 (181) | 27.2 (104) | 25.4 (97) |
| Antipseudomonal penicillin +<br>inhibitor |  |  |  |
| Piperacillin-tazobactam (CLSI 2022) | 52.9 (202) | 24.3 (93) | 22.8 (87) |
| Piperacillin-tazobactam (CLSI 2023) | 52.9 (202) | 11.0 (42) | 35.3 (138) |
| Monobactam |  |  |  |
| Aztreonam | 18.3 (70) | 29.8 (114) | 51.8 (198) |
| Antipseudomonal fluoroquinolones |  |  |  |
| Levofloxacin | 27.5 (105) | 20.7 (79) | 51.8 (198) |
| Ciprofloxacin | 26.4 (101) | 35.6 (136) | 38.0 (145) |
| Aminoglycosides |  |  |  |
| Gentamycin (CLSI 2022) | 63.4 (242) | 22.3 (85) | 14.4 (55) |
| Tobramycin (CLSI 2022) | 88.7 (339) | 2.1 (8) | 9.2 (35) |
| Tobramycin (CLSI 2023) | 51.6 (197) | 31.7 (121) | 16.8 (64) |
| Amikacin (CLSI 2022) | 88.0 (336) | 4.2 (16) | 7.1 (27) |
| Amikacin (CLSI 2023)<sup>b</sup> | 75.0 (54) | 12.5 (9) | 12.5 (9) |
| Polymyxin |  |  |  |
| Colistin | - | 99.2 (379) | 0.8 (3) |

<sup>a</sup> Unless mentioned, criteria for S/I/R are identical between CLSI M100-Ed32 (2022) and M100-Ed33 (2023).
<sup>b</sup> Amikacin MIC against urine-associated isolates only ( $n = 72$ , linked to the source categories: urine, urine collected by catheter, urine obtained from indwelling catheter, catheterized urine, and urethral discharge in Table S1).

Susceptibility data for individual isolates are provided in Table S1. Among the CRPA isolates, 27.7% (106/382) were deemed difficult-to-treat resistance *P. aeruginosa* (DTR-*P. aeruginosa*) in Tamma *et al*.’s definition (24), and 85.8% (328/382) were MDR in Magiorakos *et al*.’s definition (25). The susceptibility profiles of DTR strains are summarized in Table S2. Based on the CLSI M100-ed 32 breakpoints, tobramycin (80.2%) exhibited the highest susceptibility rate of DTR isolates among the three aminoglycosides analyzed. Only three CRPA isolates were resistant to colistin; two of the three colistin-resistant isolates (Table S1) were however DTR-*P. aeruginosa* and elicited resistance to at least one agent from each of the seven antimicrobial agent categories listed in Table 1.

The MLST profiles of 382 CRPA isolates were investigated in the PubMLST database (26). Including newly assigned ST (ST4091–ST4125), 148 STs were identified (Fig. 1A). The most frequently encountered clonal group (CG) was CG274 (comprising only ST274: type III secretion system effector protein genotype *exoS*^+^/*exoU^−^*, 9.7%), followed by CG235 (ST235: *exoS^−^*/*exoU*^+^, 7.6%), CG253 (ST253: *exoS^−^*/*exoU*^+^, 4.5%), and CG155 (ST155 and ST4122: *exoS*^+^/*exoU^−^*, 3.4%).

**FIG 1.**
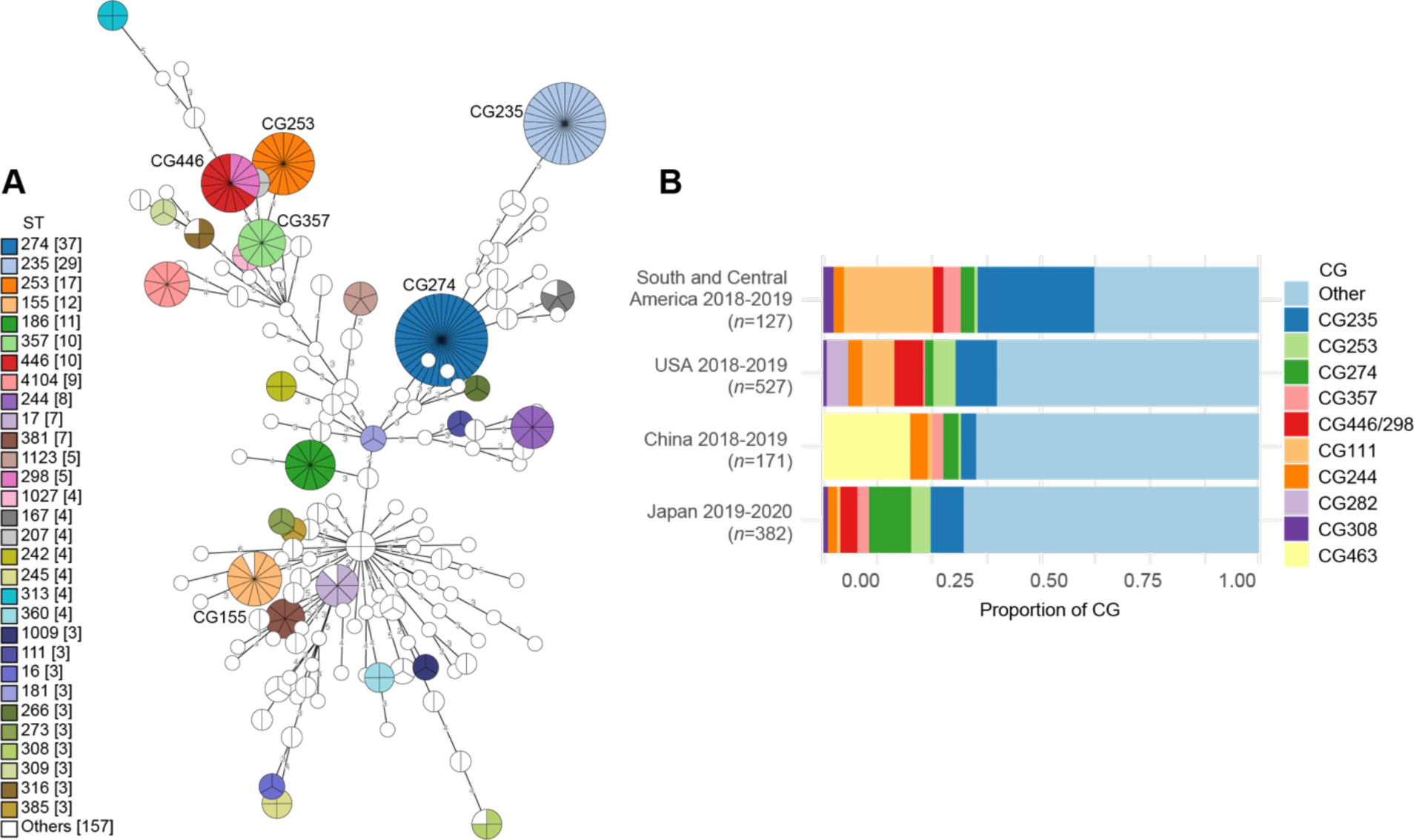
(A) Minimum spanning tree of 382 meropenem-resistant CRPA isolates based on MLST profiles. Each node represents sequence type (ST), otherwise clonal group (CG). STs with one allele difference were collapsed into one node as CG. Node size is proportional to the number of isolates. The number on the branch denotes the number of different alleles at the 7 MLST loci between two nodes. (B) Proportions of 10 globally most common CGs in Japan and other regions. Proportion of CGs in the USA, China, and South and Central America are based on Reyes *et al.*’s report (8).

Many minor STs, each contributing about 1%, occupied 49.0% of the total CRPA in Japan. We further investigated the occurrence of nearly identical isolates among different patients in our dataset by analyzing Mash distances between draft genomes (27). Among the 382 isolates examined, 7.6% (29/382) showed a Mash distance of <10^−5^ to at least one other isolate, which we defined as nearly identical. We extended this analysis to an external dataset used in Reyes *et al*’s CRPA surveillance across various regions (8). In the USA, the proportion of nearly identical CRPA isolates was 0.6% (4/527), in China it was 19.2% (33/171), in Singapore it was 4.9% (2/41), and in Central South America it was 3.5 % (6/171). Consequently, the contribution of outbreaks to the CRPA dataset in Japan is lower than in China but higher than in the USA.

Furthermore, we identified all the top ten global high-risk clones (ST111, ST175, ST233, ST235, ST244, ST277, ST298, ST308, ST357, and ST654) (28), which collectively accounted for 17.0% of the total CRPA isolates in Japan (Table S1). Reyes *et al* (8) proposed globally most common CGs (10 CGs in Fig. 1B). Those globally common CGs comprised 34.8%. Despite being among the top two most common CGs in the USA and Central South America, CG111 remains a minor CG in Japan (Fig. 1B). Surprisingly, the dominant clone CG463 (ST463), prevalent in Chinese clinical settings (7, 30), was entirely absent in the 382 CRPA isolates collected in this study. The characteristic features of two major CG/STs in Japan, ST274 and ST235 are described in more detail with genotype information later in the result section.

### Resistance mutations and carbapenemases in CRPA

The molecular mechanisms of meropenem resistance are primarily classified into four types: porin inactivation, effux pump overproduction, target alteration, and horizontal acquisition of β-lactamase (2). Those mechanisms are respectively associated with inactivation of the *oprD* gene, inactivation of either *mexR*, *nalC*, or *nalD* genes encoding transcriptional regulators (repressors) for the MexAB-OprM efflux pump (11–13), nonsynonymous substitutions in *ftsI* (14), and acquisition of plasmid or ICE-borne carbapenemase genes (4, 5). We searched for the presence of inactivating mutations (frameshift mutations due to insertion/deletion, nonsense mutations, and inactivation of start codon or stop codon) in *oprD*, *mexR*, *nalC*, and *nalD* as well as resistance mutations registered in the NCBI AMRFinderPluS database, as these likely contribute to increasing meropenem MIC.

Database searches revealed that only 4.2% (16/382) of CRPA in Japan possessed acquired carbapenemases. Known carbapenemases detected were GES-5 (*n* = 2), IMP-1 (*n* = 6), IMP-10 (*n* = 4), IMP-7 (*n* = 3), and IMP-34 (*n* = 1). No OXA-type carbapenemases were detected. ST274 CRPA isolates rarely (2.7%, 1/37) carried carbapenemase, contrasting with the 34.5% (10/29) possession rate of ST235 isolates. One ST235 isolate encoded a novel variant of IMP-type MBL, which was assigned a new allele name IMP-98 by NCBI.

The prevalence of inactivating mutations or known nonsynonymous resistance mutations in chromosomal genes in 382 CRPA isolates was 87.1% (333/382) for *oprD*, 14.1% (54/382) for *mexR*, 9.1% (35/382) for *nalD*, and 6.3% (24/382) for *nalC*. Collectively, 28.8% (110/382) of CRPA carried an inactivating mutation in at least one of the three repressors for *mexAB*. The most common mutation type in *oprD* was IN/DELs causing frameshift (237/382), followed by nonsense mutations (90/382); these include, for instance, W277stop (24/90), W339stop (9/90), and W138stop (9/90). In addition, 4.7% (18/382) of CRPA isolates carried known resistance mutations in *ftsI* (P527S, F533L, R504C, or V471G) (14).

A simultaneous occurrence of resistance mutations and/or carbapenemase was observed (Fig. 2). While 26.4% (101/382) of CRPA isolates had an inactivating mutation in both *oprD* and at least one regulator for *mexAB*, 4.5% (17/382) had both *oprD* and *ftsI* mutations, with 3.1% (12/382) exhibiting the *oprD* mutation and carbapenemase. The acquisition of carbapenemase alone occurred only in 0.7% (4/382) of the CRPA. Notably, 9.1% (35/382) did not carry identifiable inactivating mutations in the four focused genes, known *ftsI* resistance mutations, or carbapenemase.

**FIG 2.**
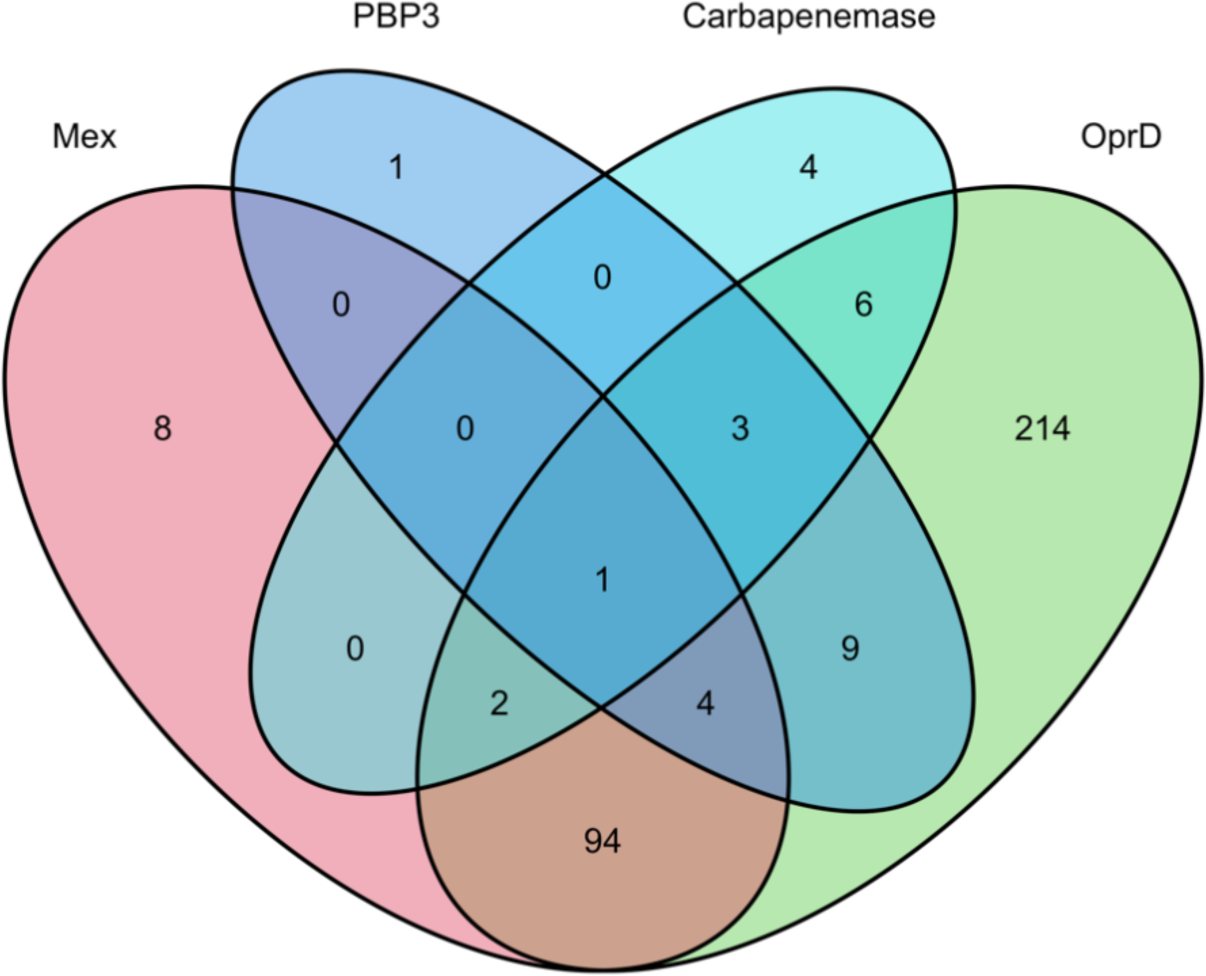
Venn diagram showing the simultaneous occurrence of resistance mutations in meropenem-resistant *P. aeruginosa*. Numbers in circles represent the numbers of isolates having an acquired carbapenemase, a point mutation registered in AMRFinderPlus (56), or inactivating mutations. The “Mex” category indicates the presence of a mutation in at least one of three loci (*mexR*, *nalC,* and *nalD*) encoding a negative regulator for the *mexAB* operon. The “PBP3” category indicates the presence of a mutation in *ftsI*. The acquired carbapenemase/MBLs detected are GES-5, IMP-1, IMP-10, IMP-7, IMP-34, and IMP-98.

### *bla*_IMP-98_

IMP-98 differed from IMP-1 by only one site at the nucleotide level and likely originated from the *bla*_IMP-1_ gene by C637T nonsynonymous substitution, leading to P213S amino acid change (Fig. 3A). To deduce the origin of the *bla*_IMP-98_ region, we determined complete genome sequence of ST235 strain, JBBCAEG-19-0032, carrying *bla*_IMP-98_ using long reads. This revealed that strain JBBCAEG-19-0032 genome consists of one chromosome, and *bla*_IMP-98_ is embedded in a class 1 integron as the upstream region of *bla*_IMP-98_ contained specific motifs of *attI1* (5ʹ-<u>GTTATGG</u>AGCAGCAACGAT<u>GTTACGC</u>AGCAGGGCAGTCG<u>CCCTAA</u>AACAAAG<u>TTAGGC</u>-3ʹ: 7-bp core sites are underlined) (29).

**FIG 3.**
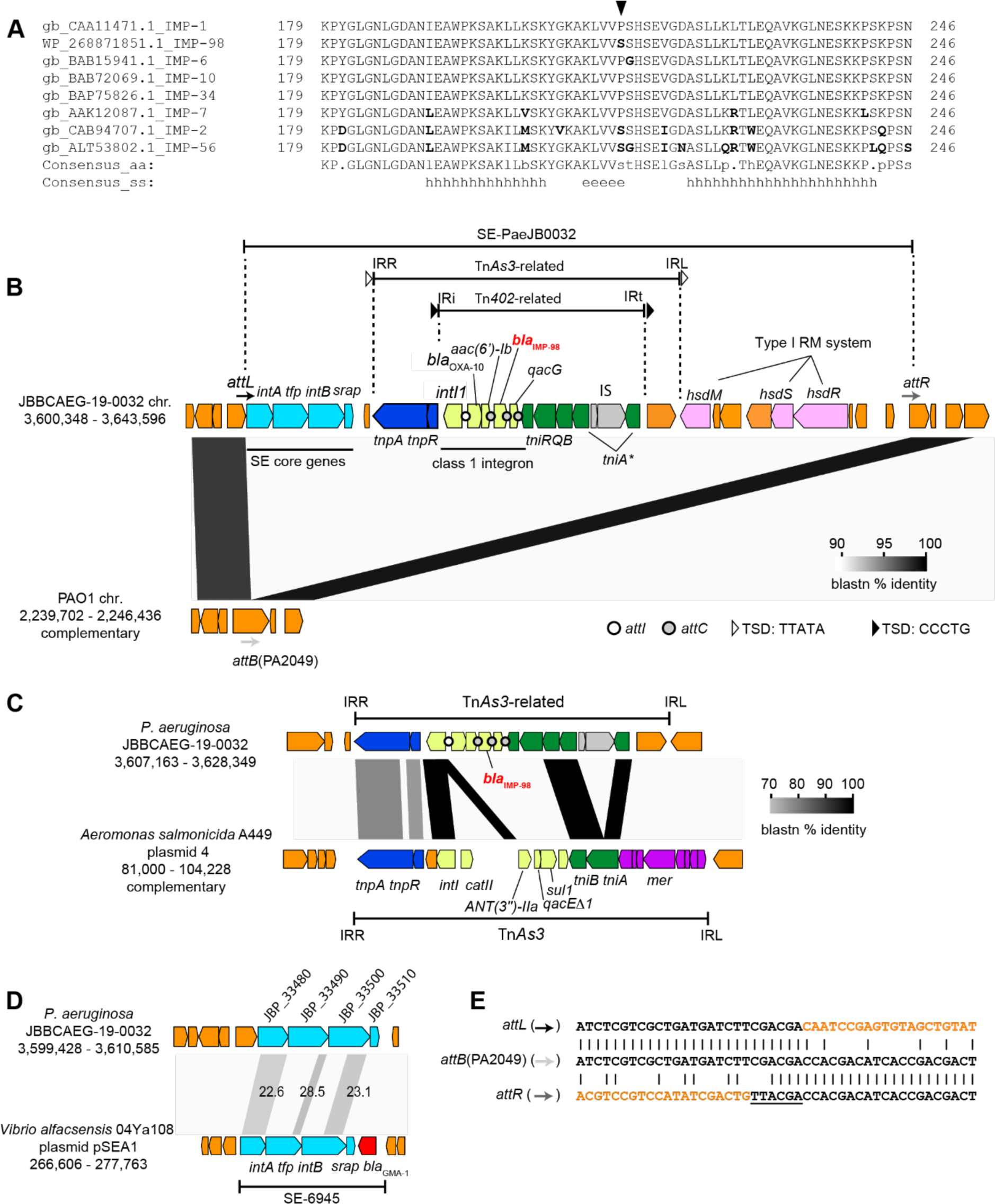
(A) Alignment of carboxyl-terminal regions of IMP variants. H, helix; e, sheet. IMP-98 differs from IMP-1 only at P213 indicated by an arrowhead. (B) Genetic context of *bla*_IMP-98_. *tni* genes in green are associated with the transposition of the Tn*402*-related element. *tniA*\* indicates disrupted *tniA*. *tnpA* and *tnpR* genes are associated with the transposition of the Tn*As3*-related transposon. The left four genes in the insertion in JBBCAEG-19-0032 are SE core genes: *intA*, encoding tyrosine recombinase; *Up*, tyrosine recombinase fold protein; *intB*, large tyrosine recombinase; *srap*, SE-associated recombination auxiliary protein (34). *bla*_OXA-10_ and *aac(6*ʹ*)-Ib* in integon overlapped by 23 bp. *intI* and *attC* were identified by IntegronFinder2.0 (59). (C) Comparison between Tn*As3* and Tn*As3*-realted elements in the JBBCAEG-19-0032 chromosome. Integron-associated genes are shown in light green. *attI* and *attC* sites were not found in Tn*As3. mer* genes in purple are mercury resistance genes. (D) Similarity of SE core genes between SE-PaeJB0032 and SE-6945 (33). The number indicates percentage identity in BLASTp. (E) Comparison of border regions between SE-PaeJB0032 and chromosome (*attL*, *attR*) with unoccupied integration site (*attB*) in PAO1. Sequences in orange are putative SE regions. The underlined sequence is a putative 6-bp footprint generated by SE integration (33, 34). Sequences are obtained from the following accession numbers: JBBCAEG-19-0032 chromosome, AP029374; PAO1 chromosome, NC_002516.2; pSEA1, AP024167; Tn*As3*, CP00654.1.

A comprehensive analysis of the *bla*_IMP-98_ context using the ISFinder database (30) revealed that the class 1 integron is situated within a Tn*402-*related transposon, which is further nested within a Tn*As3*-related transposon. The archetype Tn*As3* was originally identified in the genus *Aeromonas* (Fig. 3B) (31, 32). However, the integron structure differs between Tn*As3* and the Tn*As3*-related transposon in JBBCAEG-19-0032 in the gene cassette array composition and the absence of conserved *attC*/*attI* motifs of class 1 integron in Tn*As3* (Fig. 3C).

Both transposon units in JBBCAEG-19-0032 possess intact terminal inverted repeats and are flanked by 5-bp target site duplication (TSD): CCCTG for the Tn*402*-related transposon and TTATA for the Tn*As3*-related transposon, indicating their insertions through transposition events. Further comparison of the JBBCAEG-19-0032 chromosome with the PAO1 chromosome revealed that the Tn*As3*-related transposon is nested within a genomic island containing distant homologs of core genes of a recently defined mobile DNA element, the strand-biased circularizing integrative element (SE) (33–35) (Fig. 3BD). The products of the SE core genes in JBBCAEG-19-0032 (IntA: encoding tyrosine recombinase, Tfp: tyrosine recombinase fold protein, IntB: large tyrosine recombinase, Srap: SE-associated recombination auxiliary protein) exhibited low identities (22.6%–28.5%) in BLASTp comparisons to homologs of a characterized SE, SE-6945, from the genus *Vibrio* (Fig. 3D) (33). However, IntA and IntB homologs of JBBCAEG-19-0032 possess the catalytic RHRY motif of the tyrosine recombinase (36)(Supplemental material Fig. S1), whereas Tfp and Srap homologs share secondary structures with known *Vibrio* homologs (Fig. S2 and Fig. S3). The *in silico* removal of a 37,833-bp segment from the JBBCAEG-19-0032 chromosome, with a putative 6-bp footprint normally formed on the right side of SE after SE integration (33–35), regenerated an intact form of the PA2049-equivalent gene encoding a hypothetical protein (Fig. 3E). This suggests that the 38-kb region was inserted via site-specific recombination. Hence, we designated the 37.8-kb insert as putative SE, SE-PaeJB0032. A BLASTn search against the NCBI-nr/nt database (2024-02-07) and a database containing 382 Japanese CRPA strains using the left end (*intA* region) of the SE as a query did not yield significant hits, indicating that SE-PaeJB0032 may have recently integrated into the *P. aeruginosa* genome.

To test whether IMP-98 is functional, *bla*_IMP-98_ was cloned and introduced into the drug-susceptible *E. coli* strain DH5α, after which the susceptibility for various β-lactams was investigated (Table 2). MICs of β-lactams except for aztreonam were considerably higher against *E. coli* DH5α carrying *bla*_IMP-98_ than the control strain, confirming that IMP-98 retains the wide substrate range of MBL.

**TABLE 2.**
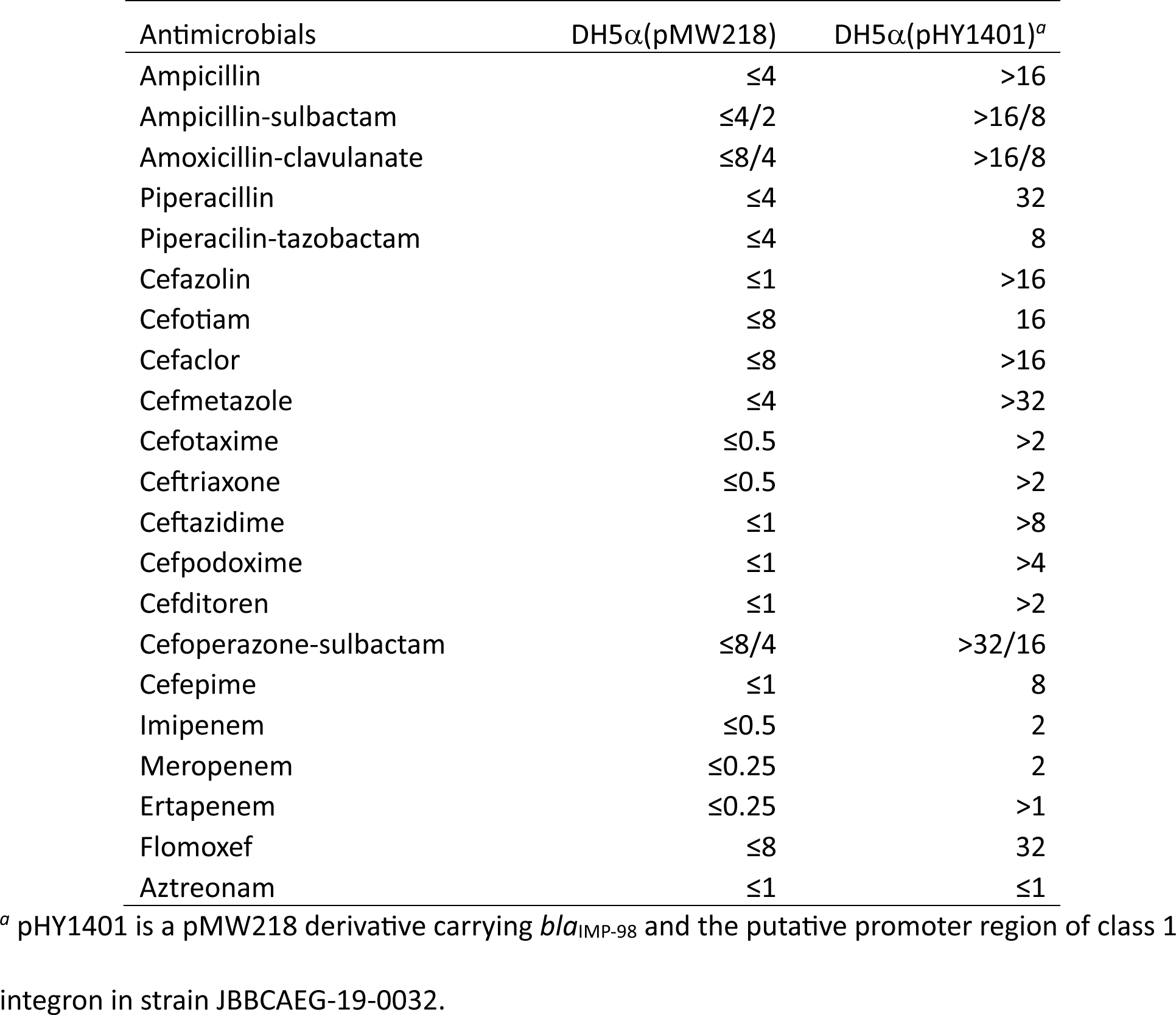
β-lactam MICs against *E. coli* strain carrying *bla*_IMP-98_.

| Antimicrobials | DH5α(pMW218) | DH5α(pHY1401)<sup>a</sup> |
| --- | --- | --- |
| Ampicillin | ≤4 | >16 |
| Ampicillin-sulbactam | ≤4/2 | >16/8 |
| Amoxicillin-clavulanate | ≤8/4 | >16/8 |
| Piperacillin | ≤4 | 32 |
| Piperacilin-tazobactam | ≤4 | 8 |
| Cefazolin | ≤1 | >16 |
| Cefotiam | ≤8 | 16 |
| Cefaclor | ≤8 | >16 |
| Cefmetazole | ≤4 | >32 |
| Cefotaxime | ≤0.5 | >2 |
| Ceftriaxone | ≤0.5 | >2 |
| Ceftazidime | ≤1 | >8 |
| Cefpodoxime | ≤1 | >4 |
| Cefditoren | ≤1 | >2 |
| Cefoperazone-sulbactam | ≤8/4 | >32/16 |
| Cefepime | ≤1 | 8 |
| Imipenem | ≤0.5 | 2 |
| Meropenem | ≤0.25 | 2 |
| Ertapenem | ≤0.25 | >1 |
| Flomoxef | ≤8 | 32 |
| Aztreonam | ≤1 | ≤1 |

<sup>a</sup> pHY1401 is a pMW218 derivative carrying *bla*<sub>IMP-98</sub> and the putative promoter region of class 1 integron in strain JBBCAEG-19-0032.

### Other resistance determinants

Among 46 amikacin-nonsusceptible CRPA isolates, 54.3% (25/46) possessed known amikacin resistance genes *aac(6’)* variants, resistance mutations such as *fusA1* (Q455R) and *amgS* (V121G) (37), or inactivating mutation in *mexZ* encoding a repressor for the MexXY effux pump contributing to aminoglycoside resistance (38) (Table S1). However, 45.7% (21/46) of amikacin-nonsusceptible CRPA did not possess identifiable amikacin resistance genes or mutations. Among 281 ciprofloxacin nonsusceptible CRPA isolates, 76.9% (216/281) possessed at least one of the quinolone resistance gene *crpP* (39) (164/281) and resistance mutations in *gyrA* (T83I, D87N, or D87G; 122/281), *gyrB* (E468D, S466F, or S466Y; 16/281), or *parC* (S87L, S87W; 46/281), *parE* (A473V; 6/281) (Table S1). Three colistin-resistant isolates did not possess any known colistin resistance mutations or genes.

### Features of two major STs in Japan

Specifically, we aim to determine whether notable phenotypic or genotypic features are associated with ST274 (*n* = 37) and ST235 (*n* = 29). Table 3 presents the observation frequencies of four source groups in ST274 and ST235, as well as other STs. The relative frequencies among the three ST groups differed significantly (*P* = 2.5 x 10^−2^, chi-square test for Table 3, df = 6). The proportion of isolates liked to the Oral/Endotracheal/Respiratory category was higher in ST274 (64.8%, 24/37) compared to the rest (48.1%, 166/345), however, this difference was not significant (*P* = 7.8 x 10^−2^, chi-square test, df = 1). A significant difference was observed in the proportion of isolates in the Urinary/Genital category between ST235 (41.3%, 12/29) and the rest (18.1%, 64/353) (*P* = 5.6 x 10^−3^), suggesting that ST235 may have a competitive advantage over other STs in urine.

**Table 3.** Isolation sources observed in two major STs.

| Source category group | ST274 | ST235 | Other STs |
| --- | --- | --- | --- |
| Oral/Endotracheal/Respiratory | 23 | 9 | 157 |
| Urinary/Genital | 4 | 12 | 60 |
| Blood/Fluids | 1 | 3 | 36 |
| Other sources | 9 | 5 | 63 |
| Total | 37 | 29 | 316 |

<sup>a</sup> More specific source categories are shown in Table S1.

Table 4 provides a summary of resistance phenotypes, genes, and mutations observed in the two major STs. Regarding nonsusceptibility, different proportions were observed among the three ST groups for piperacillin-tazobactam, ceftadizime, ciprofloxacin, and amikacin (Table 3).

**Table 4.** Observation frequency of specific resistance phenotype, genes, or mutations in two major ST groups.

| Phenotype or genotype | Percentage of relevant isolates in ST<br>(number of isolates) |  |  | Adjusted *P*-value<sup>a</sup> |
| --- | --- | --- | --- | --- |
|  | ST274 | ST235 | Other STs |  |
|  | (total 37) | (total 29) | (total 316) |  |
| Non-susceptibilities<sup>b</sup> |  |  |  |  |
| Piperacillin-tazobactam | 70.2 (26) | 58.6 (17) | 45.9 (145) | 0.02309 |
| Ceftazidime | 32.4 (12) | 65.5 (19) | 44.3 (140) | 0.04234 |
| Ciprofloxacin | 70.2 (26) | 93.1 (27) | 72.1 (228) | 0.05972 |
| Amikacin | 13.5 (5) | 62.0 (18) | 7.2 (23) | 0.00035 |
| DTR | 32.4 (12) | 48.2 (14) | 25.3 (80) | 0.04235 |
| Resistance genes/mutations |  |  |  |  |
| Carbapenem |  |  |  |  |
| *bla*<sub>IMP</sub>, or *bla*<sub>GES</sub> | 2.7 (1) | 34.5 (10) | 1.6 (5) | 0.00035 |
| *oprD*<sup>c</sup> | 89.1 (33) | 96.6 (28) | 86.1 (272) | 0.35682 |
| *mexR*, *nalC*, or *nalD*<sup>c</sup> | 27.0 (10) | 27.6 (8) | 28.4 (90) | 1.0 |
| *ftsI*<sup>c</sup> | 5.4 (2) | 10.3 (3) | 4.1 (13) | 0.37338 |
| Fluoroquinolone |  |  |  |  |
| *gyrA*, *gryB*, *parD*, or *parC*<sup>c</sup> | 59.5 (22) | 93.1 (27) | 28.5 (90) | 0.00035 |
| *crpP* | 32.4 (12) | 37.9 (11) | 63.0 (199) | 0.00084 |
| Amikacin |  |  |  |  |
| *acc(6′)-1b*, *aac(6′)-II*,<br>*aac(6′)-31*, *aacA8*,<br>*aac(6′)-Ia*, *aac(6′)-Ib3*,<br>*aph(3′)-XV*, or *aac(6′)-Iae* | 0 (0) | 58.6 (17) | 2.2 (7) | 0.00035 |

<sup>a</sup> Chi-square test *P*-value after multiple testing corrections (false discovery rate) by Benjamini &
Hochberg method. The independence of two factors (phenotype/genotype and ST) was
evaluated by using a 2 by 3 contingency table (df = 2) where the rows are the count of qualified
isolates and the count of nonqualified isolates. DTR: difficult-to-treat resistance.
<sup>b</sup> Breakpoint criteria follows CLSI M100-ed 32 (2022).
<sup>c</sup> Acquired mutation in conserved genes.

Notably, the proportion of amikacin-nonsusceptible isolates differed significantly between ST235 (62.0%, 18/29) and the rest (7.9%, 28/353) (*P* < 2.2 x 10^−16^, chi-square test, df = 1).

The proportion of DTR isolates also varied among STs (*P* = 4.2 x 10^−2^, adjusted P-value in Table 3). Particularly, the proportion of DTR isolates in ST235 (48.0%, 14/29) significantly differed from the rest (26.1%, 92/352) (*P* = 1.8 x 10^−2^). This trend was not significant in ST274 (32.4% in ST274 vs 27.2% in other STs) (*P* = 0.6).

The possession rates of chromosomal mutations in *oprD*, *mexAB* regulator genes, and *ftsI* did not differ among STs (Table 4). However, the possession rates of carbapenemase genes *bla*_IMP_ and *bla*_GES_ differed significantly among STs. The possession rates of *bla*_IMP_ and *bla*_GES_ in ST235 were significantly higher than that in other STs: 34.5% in ST235 vs 1.7% in other STs (*P* = 1.4 x 10^−15^). This trend was also observed for the possession rate of amikacin resistance genes: 58.6% in ST235 vs 2.0% in other STs (*P* < 2.2 x 10^−16^). These characteristic features of ST235 were absent in ST274.

Fluoroquinolone resistance determinants can be categorized into chromosomal resistance mutations (*gyrA*, *gryB*, *parC*, *parD*), and transferrable quinolone resistance gene *crpP* (39). Although both types appeared at different proportions among STs (Table 4), the distribution trend differed between the two types. The proportion of isolates carrying *gyr* or *par* mutations was higher in the two major STs (75.4%, 49/65) than in others (28.5%, 90/316) (*P* = 5.6 x 10^−15^), while the proportion of isolates carrying *crpP* was higher in nonST274 and nonST235 (63.0%, 199/316) than the two STs (34.8%, 23/66) (*P* = 4.6 x 10^−5^). Surprisingly, *crpP* was present in 57% (58/101) of ciprofloxacin-susceptible isolates. Thus, its significance should be interpreted with caution. These observations validate the notion that ST235 exhibits an extraordinary resistance phenotype among STs, functioning as an epidemic high-risk clone (28).

## DISCUSSION

Since the adoption of a global action plan on antimicrobial resistance at the WHO conference, regional surveillance on antimicrobial-resistant bacterial pathogens has commenced worldwide (8, 15–17, 23). However, in the case of CRPA, most surveillance projects have not provided both MIC values of antimicrobial agents and genome sequences for the collected strains simultaneously, limiting data interpretation and reuse. Furthermore, comprehensive surveillance in a single country with a 10% prevalence rate of CRPA, such as Japan, has been lacking until this study. This study was aimed at addressing these gaps.

The ciprofloxacin and amikacin susceptibility rates regarding CRPA were 26.4% and 88.0%, respectively, in Japan. These values exceed 20.6% and 60.6% of pooled CRPA values reported in the Asia–Pacific region (CLSI breakpoint criteria before 2023) (16), and are then equivalent to and higher than 26.2% and 63.4% of the global CRPA population, respectively (17).

Furthermore, the piperacillin-tazobactam susceptibility rate in Japan was 52.9%, which contrasts with the 25.3% susceptibility rate in the global CRPA population (17). This difference might reflect the difference in the carbapenemase possession rate in CRPA between Japan and other regions. Therefore, in Japan, MDR has not yet heavily prevailed in the *P. aeruginosa* population in clinical settings compared with other Asia–Pacific regions or the world average. Notably, fluoroquinolone resistance among CRPA is a global problem and is common even in non-high-risk clones.

Since several countries/regions normally categorized in the Asia–Pacific region are geographically isolated by the presence of sea, we expected to observe distinct features of the CRPA genotype in Japan and other Asia–Pacific regions. The striking feature was the predominance of CG463/ST463 in China (8) and its absence in Japan (Fig. 1B). Among the carbapenemase types, IMP was the most frequent (87.5%; 14/16) in Japan. This trend contrasts with the predominance of NDM in *P. aeruginosa* borne-carbapenemases in India (16), KPC in China (8), and VIM in Europe, Africa/Middle East, and Latin America (17). As a country with an average prevalence rate of meropenem-resistant CRPA, Japan was expected to have a low carbapenemase possession rate among *P. aeruginosa* strains. This study reveals that the carbapenemase possession rate in Japan is 4.2%, which is among the lowest compared to other regions where surveillance studies have been conducted. For instance, the carbapenemase possession rate of meropenem-resistant CRPA was 32.9% in the Asia–Pacific region from 2015 to 2019 (16), 1.9% for the USA in 2018–2019 (8), 69% for South and Central America in 2018–2019 (8), 32% for China in 2018–2019 (8), 57% in Singapore–Australia in 2018–2019 (8), and 21.9% for Europe in 2017–2019 (17).

The prevalence rate of carbapenemase is influenced by genetic, demographic, and environmental factors. The genetic factors include the type of STs constituting the *P. aeruginosa* population. As shown in Table 5, only so-called high-risk clones, represented by ST235, seem to have a trend toward capturing transferrable carbapenemase. This could be associated with the number of integrons possessed by the STs, as most reported MBL genes other than NDM are present as integron gene cassettes (28). In Japan, the proportion of high-risk clones is 17%, which is lower than that in China (30%, including KPC-producing ST463) and Singapore (60% in Reyes’s surveillance (8)). Therefore, potential carbapenemase gene recipients might be fewer in Japan than in other regions. The demographic factor includes the low migration rate of carbapenemase-producing CRPA to hospitals in Japan from high-carbapenemase prevalence countries/regions. However, this factor is challenging to evaluate based on publicly available information currently. Environmental factors include the hospital’s environmental condition that can prevent outbreaks and the resulting clonal expansion of carbapenemase-producing CRPA within the country. The low occurrence of outbreaks is supported by the low frequency (7.6%) of nearly identical isolates in the 382 CRPA dataset, as opposed to 19.2% in the Chinese CRPA dataset. Previously, VIM-producing CRPA isolates were found in hospitals in Japan (40–42); however, they were not detected in the 78 hospitals participating in the JARBS from 2019 to 2020. Furthermore, according to the National Epidemiological Surveillance of Infectious Diseases by the Ministry of Health, Labor, and Welfare, the number of amikacin-nonsusceptible, carbapenem, and fluoroquinolone-resistant *P. aeruginosa* has been decreasing over a decade in Japan (43). Therefore, it is speculated that hospital environments are being managed to reduce outbreaks and horizontal transmission of carbapenemases in Japan.

The complete sequence determination of a *bla*_IMP-98_-carrying strain revealed that the *bla*-containing-integron was embedded in a Tn*3*-family transposon of *Aeromonas* origin and nested in a putative SE region in the chromosome, rather than in the plasmid. Previous studies have identified *Pseudomonas* as one of the top five representative host genera of SEs; however, to date, only one terminus-delineated SE from *P. aeruginosa*, SE-PaeBT2436, carrying the *tmexCD-topJ* cluster conferring tetracycline and tigecycline resistance (34). Since homologs of SE core genes could not be detected by nucleotide-based search in either the current NCBI database or the CRPA dataset, the SE-PaeJB0032 type might have formed only recently, potentially capturing *bla*_IMP-98_, in other taxa such as *Aeromonas*, and then jumping into the chromosome of unique ST235 clone.

This surveillance revealed that two STs, ST274 and ST235, members of the top 10 globally most common CG in CRPA, were also the most common in Japan. ST274 and CG274 members have frequently been detected in clinical settings worldwide (44) and are known for chronic infection and intrabody evolution involving MDR in patients with cystic fibrosis in Europe (45, 46). Whether the ST274 isolates collected in this study are associated with a chronic lung infection remains unknown, but the “Oral/Endotracheal/Respiratory” source category was most frequent in ST274 (Table 4). However, the proportion of this category did not significantly differ from other STs (*P* = 0.078). Conversely, the other major ST, ST235, has more striking features than ST274 in isolation source, nonsusceptibility, and resistance gene possession rate (Table 3, Table 4). ST235 was proposed as an epidemic high-risk clone due to its worldwide occurrence with an extremely drug-resistant phenotype (28, 44). However, statistical data supporting this notion in a single surveillance has been lacking. Our data demonstrate that ST235 has a trend to acquire resistance genes and resistance phenotypes more frequently than other STs. The association of ST235 with urine is a novel finding, but the underlying mechanisms remain to be investigated.

Inactivating mutations in *oprD* were the most frequent meropenem resistance mechanism (found in 87.1%), as noted in previous genome surveillance performed in other regions, where 69% of meropenem-resistant CRPA possessed *oprD*-inactivating mutations and 22% possessed acquired carbapenemase (8). We show that in Japan, not only *oprD*-inactivating mutations but also efflux pump regulator inactivation and *ftsI* resistance mutations are more common carbapenem resistance mechanisms than the horizontal acquisition of carbapenemases. Therefore, future surveillance of *P. aeruginosa* should prioritize the study of chromosomal mutations.

Surprisingly, 9.2% (35/382) of the CRPA isolates did not possess apparent inactivating mutations, known *ftsI* resistance mutations, or carbapenemase. A recent multicenter study (9) reported that 14% of modified carbapenemase-inactivation method (mCIM)-positive CRPA, constituting 4.6% of the total examined CRPA, did not possess known carbapenemases.

Therefore, a fraction of the 9.2% of Japanese CRPA without a known resistance gene or mutation may harbor enzymes capable of partially degrading meropenem. The potentially relevant enzymes include variants of β-lactamase such as PDC and OXA (detected variants are listed in Table S1), as well as Zn^2+^-dependent imipenemase (47). However, pinpointing the causal loci of meropenem resistance in the 9.2% of Japanese CRPA is challenging with current knowledge, as these variants have been poorly characterized to date. Furthermore, known resistance mutations could not account for the mechanisms of amikacin nonsusceptibility in 45.7% of amikacin-nonsusceptible isolates. These CRPA isolates must possess unreported resistance genes or mutations in their genomes. A comparative genomics approach is required to clarify this claim in the future to fully understand and control CRPA spread.

Limitations of this research include the inability to evaluate whether CRPA isolates are associated with colonization or infection (the cause of disease) in patients when collecting isolates from hospitals, and the lack of investigation into the effectiveness of new antimicrobial agents such as cefiderocol, ceftolozane-tazobactam, and ceftazidime-avibactam against CRPA isolates carrying chromosomal resistance mutations and carbapenemases. Further surveillance studies will require new study designs to address these limitations.

## Conclusions

The first nationwide surveillance for CRPA in Japan demonstrates Japan to be a low-prevalence country for carbapenemase-producing CRPA. Although the CRPA largely consists of many minor STs in Japan, the presence of globally common CG/ST and all top 10 high-risk clones was noted. This underscores the need for continued and comprehensive surveillance. A high-risk clone ST235 has a distinguished feature among STs to become DTR relatively easily and capture carbapenemase and aminoglycoside resistance genes. The carbapenem resistance mechanisms of CRPA cannot be fully explained by known resistance genes and mutations. The dataset built in this study provides the national epidemiological features and serves as a foundation for future studies on global epidemiology, therapeutic options, and infection control.

## MATERIALS AND METHODS

### Strains, media, and plasmids

CRPA candidates participating in our genome surveillance program (the Japan Antimicrobial-Resistant Bacterial Surveillance [JARBS]) and associated with isolation source and date information were transferred from hospitals to the NIID AMR center. As part of this program, hospitals were requested to transfer all isolates showing an imipenem or meropenem MIC ≥8 in the initial test, limited to one isolate per patient. A total of 668 viable isolates were received at the AMR-RC between 2019 and 2020. Since the JARBS-PA project commenced in mid-2019, the dataset primarily consists of isolates identified in 2020.

Subsequently, 273 isolates exhibiting a meropenem MIC < 8 upon remeasurements at AMR-RC were excluded from the dataset to align with the definition of CRPA used in other surveillance studies (8, 16, 17). Additionally, three isolates lacked linked isolation source data, and 10 isolates with poor draft genome assembly quality (L90 > 150) were excluded. The final CRPA dataset comprises 382 isolates with linked isolation source data.

Single colony isolation was performed if necessary, using BD BBL™ BTB Lactose-Contained Agar Medium (also called Drigalski agar) (Becton, Dickinson and Company, Franklin Lakes, NJ, USA). BD BBL™ Muller–Hinton II (Cation adjusted) Broth (Becton, Dickinson and Company) was used to culture *P. aeruginosa* strains in liquid. Table S1 contains the *P. aeruginosa* strains and associated metadata.

*E. coli* strain DH5α [F^−^, Φ80d*lacZ*ΔM15, Δ(*lacZYA-argF*)U169, *deoR*, *recA*1, *endA*1, *hsdR*17(r_K_^−^, m_K_^+^), *phoA*, *supE*44, λ^−^, *thi*-1, *gyrA*96, *relA*1] was used for DNA cloning and functional analysis of *bla*_IMP-98_. DH5α was cultured using BD Difco™ LB, Lennox Broth and agar (Becton, Dickinson and Company). A gene cassette array of *bla*_OXA-10_-*acc(6*ʹ)-*bla*_IMP-98_ in *P. aeruginosa* ST235 strain JBBCAEG-19-0032 was PCR-amplified using primers blaIMP_F: 5ʹ-TCGAGCTCGGTACCCCGCTACTTGAAGTGTTGACGC-3ʹ and blaIMP_R: 5ʹ-CTCTAGAGGATCCCCTTAGTTGCTTGGTTTTGATGGTTTTTTACTTTCG-3ʹ, and KODone polymerase (TOYOBO CO., LTD., Osaka, Japan). A low-copy number vector pMW218 (*nptII*, pSC101 replicon; NIPPON GENE CO., LTD., Tokyo, Japan) was also PCR-amplified with primers pMW_mcs_F: 5ʹ-GGGTACCGAGCTCGAATTCGTA-3ʹ and pMW_mcs_R: 5ʹ-GGGGATCCTCTAGAGTCGACCT-3ʹ. The two PCR products were joined using NEBuilder (New England Biolabs, Ipswich, MA, USA). The resulting construct was named pHY1399. A segment on pHY1399 was PCR-amplified to delete the *bla*_OXA-10_-*acc(6*ʹ*)-Ib* region using 5ʹ phosphorylated primers P_Integ_del_F: 5ʹP-CATGGCACCTTCTTGGTGGCTAACG-3ʹ, and P_Integ_del_R: 5ʹP-GAGCGAACACGCAGTGATGCC-3ʹ, then the purified PCR product was ligated using Ligation High ver.2 (Toyobo) and introduced into DH5α. The resulting recombinant plasmid obtained from one transformant was named pHY1401.

### Antimicrobial susceptibility test

To obtain MICs of antipseudomonal agents, CRPA strains were propagated on Drigalski agar and incubated for 1 day at 35°C. Then, a single colony was inoculated into MicroScan Neg MIC NF-1J panel using Prompt Inoculation System D (Beckman Coulter Inc., Brea, CA, USA). Inoculated panels were incubated at 35°C for 18 h in MicroScan Walkway40 Plus (Beckman Coulter Inc.) or MicroScan Walkway DxM 1096 (Beckman Coulter Inc.). To cover a therapeutically achievable range, the antimicrobial susceptibility testing was designed to measure major antimicrobials using four dilutions for amikacin and colistin, and five dilutions for other antimicrobials.

To obtain β-lactam MIC against two *E. coli* strains DH5α (pMW218) and DH5α (pHY1401), the strains were first propagated on LB agar containing kanamycin (25 μg/ml) and incubated at 37°C overnight, then the colonies were diluted and inoculated into the Neg MIC EN-2J panel containing β-lactams listed in Table 2 and incubated in MicroScan Walkway40 Plus (Beckman Coulter Inc.). Breakpoint criteria for susceptible (S), intermediate resistance (I), and resistance (R) followed the CLSI 2022 guidelines M100-Ed32. The definition of DTR-*P. aeruginosa* followed Tamma *et al.*’s criteria (24): nonsusceptible to all the eight antimicrobials belonging to either five antipseudomonal agent categories, and is based on the following MIC criteria: piperacillin/tazobactam ≥32/4 µg/ml, ceftazidime ≥16 µg/ml, cefepime ≥16 µg/ml, meropenem ≥4 µg/ml, imipenem ≥4 µg/ml, aztreonam ≥16 µg/ml, ciprofloxacin ≥1 µg/ml, and levofloxacin ≥2 µg/ml (I or R in CLSI 2022). The definition of MDR was modified from Magiorakos *et al.*’s criteria (25) as follows: nonsusceptible (CLSI 2022) to at least one agent in at least three categories except for polymyxin listed in Table 1.

Notably, the equivalence of the Beckman MicroScan panel system to the broth dilution method per antimicrobial, and received the following 510(k) numbers from The United States Food and Drug Administration: imipenem, K162740; meropenem, K192355; doripenem, K101425; levofloxacin, K193358; ciprofloxacin, K193536; ceftazidime, K202343; piperacillin-tazobactam, K955910; amikacin, K862140; tobramycin, K862140; gentamycin, K862140; aztreonam, K863776; cefepime, K962150; and colistin, N62-160/S1. Essential agreement between the MicroScan Walk away method and the reference method for *P. aeruginosa* was reported to be 91.3%–96.8% (48–51)

### Genome sequencing

Draft genome was determined for all 382 CRPA isolates. For this purpose, *P. aeruginosa* strains were cultured in Muller–Hinton II broth overnight at 35°C. Cells were collected in 1.5 mL tubes and resuspended in a lysis solution containing lysozyme and RNaseA, then incubated at 37°C for 1 h. After adding proteinase K and SDS, the reaction mixture was incubated at 55°C overnight. Genomic DNA was purified from the cell lysates using Agencourt AMPure XP beads (Beckman Coulter Inc.) following the protocol recommended by the manufacturer. NGS libraries were constructed using Enzymatics 5× WGS fragmentation mix and WGS ligase reagents (Qiagen, Hilden, Germany), and then libraries were sequenced on HiSeq X Five platform in Macrogen Japan (Tokyo, Japan). Low-quality reads were removed from the original reads using fastp (52), after which the trimmed reads were assembled using Shovill pipeline (https://github.com/tseemann/shovill/tree/master) set to default options. Draft genome assemblies fulfilled L90 < 150.

To determine the complete genome sequence of strain JBBCAEG-19-0032, genomic DNA was extracted using the Monarch HMW DNA Extraction Kit for Tissues (New England Biolabs, Ipswitch, MA, USA). Long-read sequencing was performed on the GridION platform (Oxford Nanopore Technologies, Oxford, UK). The library was prepared using the Rapid Barcoding Kit, and sequencing was performed using the R9.4.1 flow cell (Oxford Nanopore Technologies).

Basecalling was performed using the high-accuracy base calling model of Guppy v5.0.12 (Oxford Nanopore Technologies). The complete sequence of JBBCAEG-19-0032 was obtained through long-read-only assembly using Flye v. 2.9-b1768, set to the “--nano-raw” option (53), followed by polishing with Illumina reads using Pilon v. 1.22 (54).

### Data analysis

The PubMLST ST number was assigned by searches against the PubMLST database (26). When hit ST was not present in PubMLST, MLST profiles of the isolates were submitted to PubMLST, after which new ST numbers (ST4091-ST4125) were assigned to the new MLST profiles. The relationship between MLST profiles was represented as a minimum spanning tree by GrapeTree software (55). Acquired antimicrobial resistance genes in draft genome assemblies were searched for using AMRFinderPlus (database ver. 2023-04-17.1) with the “--organism Pseudomonas_aeruginosa” option, otherwise default setting (56). The presence/absence and integrity of specific chromosomal genes (frameshift mutations, nonsense mutations, and gene fusions caused by substitutions at stop/start codon positions) were determined by tblastn searches against an in-house generated nucleotide database using blast v2.12.0 (57), and the following retrieval of hit sequences and inspection of the alignments. The RefSeq/GenBank accession numbers of blast queries used are as follows: ExoU, WP_034024595.1; ExoS, NP_252530.1; FtsI (PBP3), WP_003094139.1; MexR, NP_249115.1; NalC, NP_252410.1; NalD, NP_252264.1; OprD, NP_249649.1; MexZ, WP_003088626.1. The alignments were generated using MAFFT v 7.490 with the einsi option (58). The integron integrase gene and *attC* were searched for using IntegronFinder 2.0.2 with the default parameter setting (59). Secondary structure predictions for SE-associated proteins were conducted using PROMAL3D (60). Gene synteny was visualized using GenomeMatcher(61).

The genetic distance between two draft genomes was evaluated based on Mash distance (27) using Mash’s “mash dist” function with default parameter setting (k =21, s = 1000). Importantly, Mash distance serves as an approximation of 1-average nucleotide identity (ANI) (27). When required, we used Shovill genome assemblies of global CRPA isolates as external data (BioProject accession: PRJNA824880) (8). The Mash distance matrix of the 382 Japanese CRPA isolates and subsets from Reyes *et al*’s CRPA isolates were provided in the Zenodo repository.

### Data availability

Raw sequence reads generated in this study are deposited under DDBJ Sequence Read Archive accession number DRA017464 and DRA017994. Draft genome assemblies and assembly quality statistics are available from Zenodo repository [DOI: 10.5281/zenodo.10693593] (62). PubMLST allele profiles of 382 isolates and sequence alignments of *oprD*, *mexR*, *mexZ*, *nalC*, *nalD*, *ftsI*, and Mash distance matrices are available from Zenodo repository [DOI: 10.5281/zenodo.10693794] (63). The nucleotide sequence of *bla*_IMP-98_ is deposited under GenBank/DDBJ/EMBL accession number LC740578.1. Complete sequence of strain JBBCAEG-19-0032 is deposited under GenBank/DDBJ/EMBL accession number AP029374.

## ACKNOWLEDGMENTS

We thank all the hospitals contributing to the JARBS program. We thank Satoyo Wakai, Takahisa Ishizuka, Liansheng Yu, Mikihisa Okuda, Akira Moriya, Emi Fujimura, Koichi Shimakawa, Rumi Oki, Yoshie Taki, Mayumi Sasada, Mikako Nakazawa, Noriko Sakamoto, Elahi Shaheem, Chika Arai, Yuko Kazumi, Akemi Suzuki, and Fumiko Hamamoto at AMR-RC NIID for their support of isolate collection, susceptibility test, genomic DNA preparation, and JANIS data collection. We thank Enago [https://www.enago.jp] for language editing. The computation is supported by supercomputer SHIROKANE of Human Genome Center at the Institute of Medical Science University of Tokyo, and NIG supercomputer at the Research Organization of Information and Systems National Institute of Genetics. This research is supported by Japan Agency for Medical Research and Development under grant number 23fk0108604 (to M.S.). The funder has no role in study design, data collection, data analysis, and decision to publish results.

Author contributions are as follows: Conceptualization, H.Y., W.H., K.Y., and M.S.; Investigation, H.Y., W.H., Sa.K., S.A., E.A., H.Z., Sh.K., N.K., A.H., T.K., Y.S., and K.Y.; Formal analysis, H.Y., H.Z., and K.Y.; Data Curation, H.Y., N.K., and K.Y.; Methodology, H.Z., Y.S.; Visualization, H.Y.; Writing – Original Draft, H.Y.; Writing – Review & Editing, H.Y., W.H., N.K., A.H., S.K., H.Z., Y.S., K.Y., and M.S.; Project Administration and Funding Acquisition, M.S.. All authors reviewed and approved the final manuscript.

## ETHICAL STATEMENT

This study (JARBS) was reviewed and approved by the Institutional Review Board of the National Institute of Infectious Diseases (approval number 1251). The use of JANIS data was approved by Ministry of Health, Labor and Welfare of Japan (approval number 1553).

## SUPPLEMENTAL MATERIAL

**Table S1**. Susceptibility profile, genotype, and isolation source of 382 CRPA isolates.

**Table S2**. Summary of susceptibility profile of DTR isolates.

**FIG S1**. Secondary structure alignment of IntA (JBP_33480 product) and IntB (JBP_33500 product) of JBBCAEG-19-0032 with known tyrosine recombinases. R-H-R-Y motif was highlighted by yellow background. The alignment was generated using PROMAL3D. e: sheet; h, helix.

**FIG S2**. Secondary structure alignment of JBP_33490 product of JBBCAEG-19-0032 with known Tfp homologs. The alignment was generated using PROMAL3D. e: sheet; h, helix.

**FIG S3**. Secondary structure alignment of JBP_33510 of JBBCAEG-19-0032 with known Srap homologs. The alignment was generated using PROMAL3D. e: sheet; h, helix.

