## Supplementary material for "Nationwide genome surveillance of carbapenem-resistant *Pseudomonas aeruginosa* in Japan": Table S2

**TABLE S2.** Antimicrobial susceptibilities of 106 DTR-PA isolates.

| Antimicrobial agents | Percentage of susceptibility group (number of isolates)^a^ | | |
| --- | --- | --- | --- |
|  | S | I | R |
| Antipseudomonal carbapenems |  |  |  |
| Imipenem | 0 (0) | 1.9 (2) | 98.1 (104) |
| Meropenem | 0.0 (0) | 0.0 (0) | 100 (106) |
| Doripenem | 0.9 (1) | 16.0 (17) | 83.0 (88) |
| Antipseudomonal cephalosporins |  |  |  |
| Ceftazidime | 0.0 (0) | 23.6 (25) | 76.4 (81) |
| Cefepime | 0.0 (0) | 27.2 (47) | 55.7 (59) |
| Antipseudomonal penicillin + inhibitor |  |  |  |
| Piperacillin-tazobactam (CLSI 2022) | 0.0 (0) | 43.4 (46) | 56.6 (60) |
| Piperacillin-tazobactam (CLSI 2023) | 0.0 (0) | 16.0 (17) | 84.0 (89) |
| Monobactam |  |  |  |
| Aztreonam | 0.0 (0) | 22.6 (24) | 77.4 (82) |
| Antipseudomonal fluoroquinolones |  |  |  |
| Levofloxacin | 0.0 (0) | 17.9 (19) | 82.1 (87) |
| Ciprofloxacin | 0.0 (0) | 32.1 (34) | 67.9 (72) |
| Aminoglycosides |  |  |  |
| Gentamycin (CLSI 2022) | 42.4 (45) | 31.1 (33) | 26.4 (28) |
| Tobramycin (CLSI 2022) | 80.2 (85) | 2.8 (3) | 17.0 (18) |
| Tobramycin (CLSI 2023) | 39.6 (42) | 29.2 (31) | 31.1 (33) |
| Amikacin (CLSI 2022) | 76.4 (81) | 9.4 (10) | 14.1 (15) |
| Amikacin (CLSI 2023)^b^ | 56.0 (14) | 20.0 (5) | 24.0 (6) |
| Polymyxin |  |  |  |
| Colistin | - | 98.1 (104) | 1.9 (2) |
| a Unless mentioned, criteria for S/I/R are identical between CLSI M100-Ed32 (2022) and M100-Ed33 (2023). | | | |
| ^b^ Amikacin MIC against urine-associated isolates only (n=25) | | |  |
