## Supplementary material for "Nationwide genome surveillance of carbapenem-resistant *Pseudomonas aeruginosa* in Japan": Fig. S1

```
Conservation:                  9
IntA_JBBCAEG-19-0032        1  MARNH----------LRSKTG-QGR---YES--------------KARRVLIPEFRLLR-INFLE-----   36
IntB_JBBCAEG-19-0032        1  MSIIAH---------LQFSQVQDHQLPDRCNSAS-------QYLN-VDDNFVISRTAQG-VIGSIYGDDE   52
Int_lambda_NP_040609.1      1  MGR-------------------------RRS--------------HERRDLPPNLYIRN-NGYY------   24
IntI1_R388_HXF36_RS00185    1  MKTAT-----------------------------------------------------------------    5
IntB_6945_BBM67682.1        1  MYSIS--HDHA----LFKK---------YALPI-------DEAYPMPTARTVVSVKRDG-SPASYFEDDA   47
IntA_6945_BBM67680.1        1  MEHFYH---------LVAT---------------------SRSIEI----GDIELGDPP-GSSDFT----   31
IntB_6283_BCN27153.1        1  MIKISDLEHQNKL--LKAQQN-IGAHTNYLIDVLKPAIERGDWDELE--ALDIGFTVTGESLGKIADDEA   65
XerD_Eco_IEU92_RS14660      1  MKQ--DLARIEQ----------------------------------------------------------   10
Int_SXT_AAL59748.1          1  MAL-SVSW-------LDARLN-KEA--------------KETVVKADRDGLSARVSPK------------   35
IntA_6283_BCN27155.1        1  MIGTHEKQSFDIYDDYWLLSIEVDTLAKLQNYA---------NIDI--DTGEVTHSPYLSP---------   50
Consensus_aa:                  M....................................................ph...............
Consensus_ss:                                 hhh                                 eeeee


Conservation:                                                                       5
IntA_JBBCAEG-19-0032       37  -----THNDGNPKGLPDYE----------------------------FSSAEDSYLPNFPVLIQGH----   69
IntB_JBBCAEG-19-0032       53  WNVKM--YDAAGICVYNFSAWTEGITTPLVTVIAGEMKKIQIARMYLFQKPRKPNS----LRLGD-----  111
Int_lambda_NP_040609.1     25  ----------------------------------------------CYRDPRTGKE----FGLGRD----   40
IntI1_R388_HXF36_RS00185       ----------------------------------------------------------------------
IntB_6945_BBM67682.1       48  WDFND-LFNTKNEKKSNYT----------------------------LTF--QAQKHN------------   74
IntA_6945_BBM67680.1       32  ------VSKPMHFDGVNLL----------------------------FTS--NGE---------------   50
IntB_6283_BCN27153.1       66  WRNLQSHIEPQPKKFYTLE----------------------------FTN--DGRVVE------------   93
XerD_Eco_IEU92_RS14660         ----------------------------------------------------------------------
Int_SXT_AAL59748.1         36  -----------GKIVFQFR----------------------------YRF--DGKQQR--VDIGTYPLMK   62
IntA_6283_BCN27155.1       51  ------SNLKSRLKKTDII----------------------------IAP--DGT---------------   69
Consensus_aa:                  ...............................................hp...ptp...............
Consensus_ss:                                   ee                            e


Conservation:
IntA_JBBCAEG-19-0032       70  ----------------------------------------------GEPWAIGNLYLTTKLQREAG----   89
IntB_JBBCAEG-19-0032      112  LNKLARLALNNNLPLSDLFDDANLNRLLLPS-----FAE-LDRNPMRGILQLLKELFDIRVKHPDFK---  172
Int_lambda_NP_040609.1     41  RRIAITEAI-----------QANIELFSGHKH-KPLTAR-INSDNSVTLHSWLDRYEKILASRG------   91
IntI1_R388_HXF36_RS00185    6  -----------------------------------------APLPPLRSVKVLDQLRERIRYLH------   28
IntB_6945_BBM67682.1       75  PKLLLEFKQ----------------------------------------------RMYWLIWGGKDTLLS   98
IntA_6945_BBM67680.1       51  --PEFYANA----------------------------------------------FIMSRRIVE------   66
IntB_6283_BCN27153.1       94  RNLKNELKV---------------------------------------------LILKMMWLSPHD----  114
XerD_Eco_IEU92_RS14660     11  -------------------------------------------------------FLDALWLEKN-----   20
Int_SXT_AAL59748.1         63  LAEARNELD-----------RLRAVLDQGRNPKLYLQQERAKYSANQSFESIFRDWIDSAGKQG------  115
IntA_6283_BCN27155.1       70  ------------------------------------------------IVYPQSLYLVSKLRGE------   85
Consensus_aa:                  .......................................................hh....b........
Consensus_ss:                  hhhhhhhhh                                        hhhhhhhhhhhhh


Conservation:                               7                5
IntA_JBBCAEG-19-0032       90  ---------YESRTFRG-IADHLLDY-LRFLEDEGMD---YMHF---------------PKNDRLKVTYR  130
IntB_JBBCAEG-19-0032      173  -------FAPSDYTIIECMQALYNKHPKRQQHDPQQT---------------------------KLIPSR  208
Int_lambda_NP_040609.1     92  ---------IKQKTLIN-YMSKIKAI-RRGLPDAPLE---------------------------DITTKE  123
IntI1_R388_HXF36_RS00185   29  ---------YSLRTEQA-YVNWVRAF-IRFHGVRHPA---------------------------TLGSSE   60
IntB_6945_BBM67682.1       99  MEGKTFRKIEQVKDIQV-HVDLLLRV-FANTANNAFS---------------------------YISNDI  139
IntA_6945_BBM67680.1       67  ----------GVKDTEP-TSYALLRF-FRYLGKNNLN---------------------------WND---   94
IntB_6283_BCN27153.1      115  ---------HSFCTLYE-ALNDLKKV-INPLLNEGINTLS------------------------ALDFDQ  149
XerD_Eco_IEU92_RS14660     21  ---------LAENTLNA-YRRDLSMM-VEWLHHRGLT-------------------------LATAQSDD   54
Int_SXT_AAL59748.1        116  ---------LKEKTWHY-QKRSSEIYLLPRLG--KYP-------------------------LTDINELS  148
IntA_6283_BCN27155.1       86  ---------AAVKDTGS-IAKGLLAF-TRYLDSTHYS---QVDEDGYEIPPEYLTYKTLSKYEEEGVPWR  141
Consensus_aa:                  .............sh...h...h..h....h.p..hs.............................s..p
Consensus_ss:                             hhhhhh hhhhhhhh hhhhhh                               h  hhh


Conservation:                      5
IntA_JBBCAEG-19-0032      131  YRQRLI----------------------------------------------------------------  136
IntB_JBBCAEG-19-0032      209  LYAALI----------------------------------------------------------------  214
Int_lambda_NP_040609.1    124  IAAMLN----------------------------------------------------------------  129
IntI1_R388_HXF36_RS00185   61  VEAFLS----------------------------------------------------------------   66
IntB_6945_BBM67682.1      140  VFSQIKESLRGWSEKSCATRLNALSVLVQVNLHFPERARFHIPYQEGETARKVSKKYAAHGKGHFPTVIP  209
IntA_6945_BBM67680.1       95  ----ENEEL----ERYP----------------------------------------------------I  104
IntB_6283_BCN27153.1      150  LETWVLTDFTDI-DFEREKIYNGLNRLYLEARGLP----FKVNLNKKLTASDFGLTLKEAK---QYTVIP  211
XerD_Eco_IEU92_RS14660     55  LQALL-----------------------------------------------------------------   59
Int_SXT_AAL59748.1        149  LRNCL-----------------------------------------------------------------  153
IntA_6283_BCN27155.1      142  FAEFLL----------------------------------------------------------------  147
Consensus_aa:                  h...l.................................................................
Consensus_ss:                  hhhhhh


Conservation:
IntA_JBBCAEG-19-0032      137  ---------------------------------DFVTAGLL--------------------SASTASARI  153
IntB_JBBCAEG-19-0032      215  ---------------------------------AALNL-----------------------ELDDFNAIA  228
Int_lambda_NP_040609.1    130  ---------------------------------GYIDEG----------------------KAASAKLIR  144
IntI1_R388_HXF36_RS00185   67  ---------------------------------WLANERKV--------------------SVSTHRQAL   83
IntB_6945_BBM67682.1      210  VI------------------YEQYL-------SRLIQDVESAYRDFLGGRDLESEVEAEYRSLQTEHDIE  254
IntA_6945_BBM67680.1      105  YL------------------FRNYL-------DKQIAKGNM--------------------RRSVGASTL  129
IntB_6283_BCN27153.1      212  QRLYYMGLQKSEEVINEAYALRHELEQLADYITTYFEK----------AYVGY--------AKYLVSGDA  263
XerD_Eco_IEU92_RS14660     60  --------------------------------AERLEGGY---------------------KATSSARLL   76
Int_SXT_AAL59748.1        154  --------------------------------REVSES-----------------------SPSNTERLV  168
IntA_6283_BCN27155.1      148  ---------------------------------ANCKNIEGS-----NGDEAL--------SLSTAKSYM  171
Consensus_aa:                  ...................................hp........................p..s.p...
Consensus_ss:                                                  hhhhhh                        hhhhhhhh


Conservation:                        6
IntA_JBBCAEG-19-0032      154  NAVVNFYRRIINWN------------------------------LVS-----DANIEGVAFTEIKKYISV  188
IntB_JBBCAEG-19-0032      229  ESIAELYKQKNENT------------------------------LFG-----IPECRSRWYKNPIVWGDA  263
Int_lambda_NP_040609.1    145  STLSDAFREAIAEG------------------------------HIT-----------------------  161
IntI1_R388_HXF36_RS00185   84  AALLFFYGKVLCTD------------------------------LPW-----------------------  100
IntB_6945_BBM67682.1      255  QEIETLYQELIVDI------------------------------EVEEKAKNSG-------------IFP  281
IntA_6945_BBM67680.1      130  SVIRRFYVFCYRHG------------------------------YVE-----------------------  146
IntB_6283_BCN27153.1      264  RQKNGGITWYLVKSGERNKKRTTAFQQAFTALRSPNEEEVLALAQLHKPQIKSE----------------  317
XerD_Eco_IEU92_RS14660     77  SAVRRLFQYLYREK------------------------------FRE-----------------------   93
Int_SXT_AAL59748.1        169  SVLHKFYDWLIDEQ------------------------------ILE-----------------------  185
IntA_6283_BCN27155.1      172  SAVIGFYKWMQKYG------------------------------YIP-----NNDQNVVTHYTTRTRHYE  206
Consensus_aa:                  p.l..h@pbhb.................................bh........................
Consensus_ss:                  hhhhhhhhhhhh


Conservation:                                                                                    5
IntA_JBBCAEG-19-0032      189  TSNYGIPR------------------------------NLMVNSHNLAIANTKKQSQAEYINDGGDL-RP  227
IntB_JBBCAEG-19-0032      264  IQQHGLAS------------------------------AFESRS-------------------------I  278
Int_lambda_NP_040609.1    162  -TNHVAATR-----------------------------AAKSEVRR----------------------SR  179
IntI1_R388_HXF36_RS00185  101  ----LQEIG-----------------------------RPRPSR---------------------RLPVV  116
IntB_6945_BBM67682.1      282  QEVLDDYRDKAKDEVESIYRSKAHRGVIDAYKKDNVLLDVKAY---------------------AQL-KG  329
IntA_6945_BBM67680.1      147  -KLPFEVHGH----NKYGQML------TDITIK-----SPKQE---------------------TEL-TP  178
IntB_6283_BCN27153.1      318  ---YIDKFH-----------------------------SERQLT-------------------IGQW-TI  335
XerD_Eco_IEU92_RS14660     94  -DDPSAHLA-----------------------------SPKLPQ--------------------RLP-KD  112
Int_SXT_AAL59748.1        186  -INAAAGIT-----------------------------AKKVSG-----------------KKGKRT-RV  207
IntA_6283_BCN27155.1      207  LNHHDM-LAHTKPGSEHSNET------SNIMRM-----FPKKDT------------TPA----YKKL-KP  247
Consensus_aa:                  .p....................................s.c........................ph.p.
Consensus_ss:


Conservation:                                                  59 9  6
IntA_JBBCAEG-19-0032      228  LTVSEQSYMLQALLES---SREYQLMFYFALFTGARVQTIGTLRIKHLLG-GLDSN------GYLRLPIG  287
IntB_JBBCAEG-19-0032      279  ENWIDLKAYLGELQ--------TAAKYWIHFFTGMRSNEARHLPADSYTT-IKVNG------ADVHILRG  333
Int_lambda_NP_040609.1    180  LTADEYLKIYQAAESS---PCWLRLAMELAVVTGQRVGDLCEMKWSDIV------D------GYLYVEQS  234
IntI1_R388_HXF36_RS00185  117  LTPDEVVRILGFLE------GEHRLFAQLLYGTGMRISEGLQLRVKDLD----FDH------GTIIVREG  170
IntB_6945_BBM67682.1      330  VTEKQAIDAFRMI--------EGACFGACSAFTGMRISELTQIEGDSYKEVD-IDG------VKLCTMRS  384
IntA_6945_BBM67680.1      179  MNDLDIRHIRDNWHRAG-ISGEFRLLVATVINSGLRAIEVADIKPRHFK----VPKGFKG--KTYTDIKI  241
IntB_6283_BCN27153.1      336  TNRTEAQSLFKMLN--------GGCLWGLMARTGMRGDEMYSLNTVQGCTKDIIDR------QNIYVIHA  391
XerD_Eco_IEU92_RS14660    113  LSEAQVERLLQAPLIDQPLELRDKAMLEVLYATGLRVSELVGLTMSDIS----LRQ-------GVVRVIG  171
Int_SXT_AAL59748.1        208  LNDNEIRILWRYLHESK-ITEKNRIYIKLLLLLGGRKGELIQAEKHHFD----LQS-------AMWTVPI  265
IntA_6283_BCN27155.1      248  MHPEHKAIFNEHATSL---PKAANLIFRLCLEAGLRIDEATHFPSSDIG----IADCSDLDVVPFKITRT  310
Consensus_aa:                  hs..c...h.p.h...........hhh.hhh.hG.R..Eh.ph..pph.....h..........h.h...
Consensus_ss:                    hhhhhhhhhhhh      hhhhhhhhhhhhh   hhhhh   hhh               eeeeeee


Conservation:                          9
IntA_JBBCAEG-19-0032      288  -ASTLVDTK-----NGKR---MTLLVPG---WVVQDMLVYCRSAEAKKRRARSYYG--------------  331
IntB_JBBCAEG-19-0032      334  -----YTSKIAGQ-NHTE---TFWMTAG---IIEKGIAAALHIGKIAALINNFDD---------------  376
Int_lambda_NP_040609.1    235  --------K-----TGVK---IAIPTAL----HIDALGISMK-----ETLDKCKE---------------  264
IntI1_R388_HXF36_RS00185  171  --------K-----GSKD---RALMLPE---SLAPSLREQLSRARAWWLKDQAEGRSGVALPDALERKYP  221
IntB_6945_BBM67682.1      385  --WTDKLDK-----LSRE---DAWACAPICEKALLILTVLNDKYRSVSG---------------------  423
IntA_6945_BBM67680.1      242  GPSHGCYTK-----YSKN---RTISMPV---WLMEQVTQYCESERYKERKRLYFFN--------------  286
IntB_6283_BCN27153.1      392  -----NLSKTVKGSQSIQDEFVTTEIGM---KAYEVLQALHTPLRKRD----------------------  431
XerD_Eco_IEU92_RS14660    172  --------K-----GNKE---RLVPLGE---EAVYWLETYLEHGRPWLL---------------------  201
Int_SXT_AAL59748.1        266  -----EIRK-----QGEK---IGAPIMR---PLIKPAIELIELAMQM-----------------------  296
IntA_6283_BCN27155.1      311  --------K-----GSKP---RIVEVPI---TLYEELEIYKESHARLKKLNKRNDLIQ------------  349
Consensus_aa:                  ........K......t.p....hh.hs.....hhb.h..h.c............................
Consensus_ss:                       e               eeee  h   hhhhhhhhhhhhhhhhhhhhhh


Conservation:                                           6
IntA_JBBCAEG-19-0032      332  ----DTEDN--------------YVF-------------------LSKN-----------GVPYYTSKKE  353
IntB_JBBCAEG-19-0032      377  ----SDPSQY-------------PLFPSLVKPRKKEIRVFAGGPVISQS-----------GRSVS-----  413
Int_lambda_NP_040609.1    265  ----ILGGE--------------TII-------------------ASTR-----------REPLS-----  281
IntI1_R388_HXF36_RS00185  222  RAGHSWPWF--------------WVF-------------------AQHTHSTDPRSGVVRRHHMY-----  253
IntB_6945_BBM67682.1      424  ---DIHLSPRFSLKGEGKGWTGDTIH-------------------RQLK-----------DAQLH-----  455
IntA_6945_BBM67680.1      287  ---TGDEDA--------------PVF-------------------ITKE-----------GNRFKEE---  306
IntB_6283_BCN27153.1      432  -----SSSQ--------------SFF-------------------HKTTEDFS----KVGKASIG-----  454
XerD_Eco_IEU92_RS14660    202  ---NGVSID--------------VLF-------------------PSQR-----------AQQMT-----  219
Int_SXT_AAL59748.1        297  -----SKST--------------YLF-------------------PANG-----------QEELA-----  312
IntA_6283_BCN27155.1      350  SGKETDTIE--------------YLF-------------------LSNK-----------GKPYS-----  370
Consensus_aa:                  ........................l@....................ppp..............h......
Consensus_ss:                                          ee                   e


Conservation:
IntA_JBBCAEG-19-0032      354  ILDRRDSRISVGAGMADRASEVSIQDGAALRTHI-------QQILLPRIH--------------------  396
IntB_JBBCAEG-19-0032      414  --------------------------QQRLLARWPDLIIQEADIRELEQ-------------FDGFRDWR  444
Int_lambda_NP_040609.1    282  --------------------------SGTVSRYF-------MRARKA-----------------------  295
IntI1_R388_HXF36_RS00185  254  --------------------------DQTFQRAF-------KRAVEQ-----------------------  267
IntB_6945_BBM67682.1      456  --------------------------TKNLRQIF-------YDYSLHIDIRYLPEEMDEIFNLLNPIVHE  492
IntA_6945_BBM67680.1      307  -----------------------GQRSNSIDTLW-------GRLRNAIK------------ENS------  328
IntB_6283_BCN27153.1      455  --------------------------KYSMAWFK-------NALREELAL--TNEDLTDL-KVSDPNLS-  487
XerD_Eco_IEU92_RS14660    220  --------------------------RQTFWHRI-------KHYAVLAGI--------------------  236
Int_SXT_AAL59748.1        313  --------------------------TNGFDTTI-------PNNVKIWAR------RSL-----------  332
IntA_6283_BCN27155.1      371  --------------------------VNTLETDF-------SKLRRRIRE--------------------  387
Consensus_aa:                  ............................sh...h........ph..........................
Consensus_ss:                                            hhhhhhhh       hhhhhhh


Conservation:                                        9 69 5 5                      5    69     6  9
IntA_JBBCAEG-19-0032      397  --------RGNPEFQ---YFTFHDLRATFGMNLLESQLQHMGDQSITAALEYVQQRMGHSNKETTMQYLS  455
IntB_JBBCAEG-19-0032      445  NDPEV----TAGVVW---PLGTHQCRRSLAVYCARSGL---------VSVGSLGLQFKHLTEVMASYYRK  498
Int_lambda_NP_040609.1    296  ----------SGLSFEGDPPTFHELR-SLSARLYEKQ----------ISDKFAQHLLGHKSDTMASQYRD  344
IntI1_R388_HXF36_RS00185  268  ----------AGITK---PATPHTLRHSFATALLRSG----------YDIRTVQDLLGHSDVSTTMIYTH  314
IntB_6945_BBM67682.1      493  KLSPIKENERGELYW---RFNTHSLRRTFAHFVVGHGL---------VSLASLKHQFKHIHLAMTAIYAS  550
IntA_6945_BBM67680.1      329  -----------NPHF---DHDFHDTRATYAANKLDLLLNI-PDLSTTQALKLLKDELGHKDLSVTMRYLT  383
IntB_6283_BCN27153.1      488  --------FELGSNY---EFTPHQLRRSFAYYLIGYEL---------CNFPQLKQQFSHVSMAMTRHYAK  537
XerD_Eco_IEU92_RS14660    237  ------------DSE---KLSPHVLRHAFATHLLNHG----------ADLRVVQMLLGHSDLSTTQIYTH  281
Int_SXT_AAL59748.1        333  ----------SVEME---HWSMHDLRRTMRTRMSAIT-----------TQEVAELMIGHSKKGLDAIYNQ  378
IntA_6283_BCN27155.1      388  ----------VDRSW---YYRIHDLRSTFATHWLWNESKE-RDVGYDYLMDELKELMGHSSTSITEKYIK  443
Consensus_aa:                  ..........ss..b....hs.HphR.ohth.hh.............hs...hp.bh.H.p.shh..Y.p
Consensus_ss:                                         hhhhhhhhhhhh              hhhhhhhh    hhhhhhhhh


Conservation:
IntA_JBBCAEG-19-0032      456  -------YKSRL----------QWKN--------------------------------------------  464
IntB_JBBCAEG-19-0032      499  GSAFAVNFLNTE----------DAKNLIDSMEYERQKAQYIDYEANVINTTSRLWGGEGNRIQVARDKGQ  558
Int_lambda_NP_040609.1    345  -------DRGRE----------------------------------------------------------  349
IntI1_R388_HXF36_RS00185  315  -------VLKVG----------------------------------------------------------  319
IntB_6945_BBM67682.1      551  -------HSEVLTLLGIQN-PAKIKEKIQEQEIE-SHAEYLKDMI---EHPEEQSGGFMKFMSGEPMV--  606
IntA_6945_BBM67680.1      384  -------HWEGN----------PTKNQV---------PELMMALL---EEEHII----------------  408
IntB_6283_BCN27153.1      538  -------NASKF--QKIRQKKKTLANEIDEERID-QKARVYLSIYEKLANKEKIAGGKGKEFAKSITKTN  597
XerD_Eco_IEU92_RS14660    282  -------V--------------------ATERLR----QLHQQHH-------PRA---------------  298
Int_SXT_AAL59748.1        379  -------YQYLD-----------------------EMRHAYDVWYQQLE---------------TIIEPT  403
IntA_6283_BCN27155.1      444  -------YMNKL----------DDQI---------------NMAKAKNL---KINGGWS-----------  467
Consensus_aa:                  ......................................................................
Consensus_ss:                         h hhh          hhhh           hhhhhhhh   h


Conservation:
IntA_JBBCAEG-19-0032           ----------------------------------------------------------------------
IntB_JBBCAEG-19-0032      559  PLIITT--DRAM-----TERKFLKGELAYKSGPIG-GCTNLEPCDRISFTSIFACI--DCEKSILDDDRS  618
Int_lambda_NP_040609.1         ----------------------------------------------------------------------
IntI1_R388_HXF36_RS00185       ----------------------------------------------------------------------
IntB_6945_BBM67682.1      607  ------------VGDARFNQLVEDTKGANKSTGFG-TCFSGELCSMGHLFEPSKCVGRDCENLNVNKVEA  663
IntA_6945_BBM67680.1           ----------------------------------------------------------------------
IntB_6283_BCN27153.1      598  RNLFTDKVDNDMLTLNFWKKQIRNKKRHLHAVAPGIYCTST-TCGLRTLINLIECV--DCKNDYI--VDA  662
XerD_Eco_IEU92_RS14660         ----------------------------------------------------------------------
Int_SXT_AAL59748.1        404  GFPFNWRFGQ------------------------------------------------------------  413
IntA_6283_BCN27155.1           ----------------------------------------------------------------------
Consensus_aa:                  ......................................................................
Consensus_ss:


Conservation:
IntA_JBBCAEG-19-0032      465  ------------------------------------SVQHEFETQLFRHVN---TSANA----GATV---  488
IntB_JBBCAEG-19-0032      619  LKK---IKRGLNNLRREQAFYVAENAQYMQLESEIAAIYE---KLEKRGLR---KKMEALA---------  670
Int_lambda_NP_040609.1    350  ----------------------------------WDKIEIK-----------------------------  356
IntI1_R388_HXF36_RS00185  320  ----------------------------------GAGVRSPLDALPPLTSER------------------  337
IntB_6945_BBM67682.1      664  ENWVIRRERCIEKIEKMKEMGMFNRSSLATQLSDIRTAEK---VMADHNIQFEKYRVEAL----------  720
IntA_6945_BBM67680.1           ----------------------------------------------------------------------
IntB_6283_BCN27153.1      663  VFAEAKRKEAEIHMLYDIENNELTPQTASESYIKVQAAER---IMSDLGIDYEPVVFPNEVRDLLIPFGV  729
XerD_Eco_IEU92_RS14660         ----------------------------------------------------------------------
Int_SXT_AAL59748.1             ----------------------------------------------------------------------
IntA_6283_BCN27155.1           ----------------------------------------------------------------------
Consensus_aa:                  ......................................................................
Consensus_ss:                                                    hhhhhh   h


Conservation:
IntA_JBBCAEG-19-0032           --
IntB_JBBCAEG-19-0032           --
Int_lambda_NP_040609.1         --
IntI1_R388_HXF36_RS00185       --
IntB_6945_BBM67682.1           --
IntA_6945_BBM67680.1           --
IntB_6283_BCN27153.1      730  FS  731
XerD_Eco_IEU92_RS14660         --
Int_SXT_AAL59748.1             --
IntA_6283_BCN27155.1           --
Consensus_aa:                  ..
Consensus_ss:
```

**FIG S1.** Secondary structure alignment of IntA (JBP_33480) and IntB (JBP_33500) of JBBCAEG-19-0032 with known tyrosine recombinases. R-H-R-Y motif was highlighted by yellow background. The alignment was generated using PROMAL3D. e: sheet; h, helix.
