## Supplementary material for "Nationwide genome surveillance of carbapenem-resistant *Pseudomonas aeruginosa* in Japan": Fig. S2

```
Conservation:                   9                             6                           66
WP_013855472.1_VagAT68554    1  MAKSRLSVSRNQNKPITQLKAKD------NVTSLDLS---FELHQRGMAYSIKYNLLDWCHE--------   53
BCN27154.1_6283              1  MAKSKIA---KQSKAVKHVRNNQ------NVVPLNLD---IPYKGTRETRVKLFDVSHLLYF--------   50
WP_017821119.1_ValAT17749    1  MTE------------LTYLSGWR----------SDLC---IKLDV---RDNLLADFNRLLFN--------   34
BCN27178.1_6945              1  MTENFDSDKPDNVRTYRSIPTDEMQSDNPLSKIHQWK---YIWVS-GNGGNRTCDFKGWLKY--------   58
JBBCAEG-19-0032_JBP_33490    1  MI-SDLV---QRS--VVIIDAKDK-----NVEIIHPQLTVLMFDNKSGAGKRHLDLGALCYVRRGRGTPQ   59
WP_014386728.1_Vda04Ya311    1  MSRK------DNSKPVVLDDAFYEAPAA-TVEPLHLDELKLK------CGGATSHFIQLLYQ--------   49
Consensus_aa:                   Mhcp......pps..l..lps.p.......sp.hphp...hbh.s...s.s.hhch..hh@.........
Consensus_ss:                                  eee            eee eee   eee        eeeehhhhhh


Conservation:                                           7                   6                  7
WP_013855472.1_VagAT68554   54  --QCDP----QK------PLVKPSRLNRMQKLRAWVNQERKNQSSEAYIRVKLASLKQYISFCDFNMLD-  110
BCN27154.1_6283             51  --GSNK----EN------KKI-GNRTVFIRSFCKKAHQYVSNGKSATSVTGCYESLRAYVAFCDALKVD-  106
WP_017821119.1_ValAT17749   35  --ELPS----SKELQQSNQFKFADREAFVFQLKDSFSEKIDEGVSHDSLYGIYVEASQYLRWCDKESEP-   97
BCN27178.1_6945             59  -----------------------STKEHVSLLISKFEQ---SGWAEASKVRNFQHIDNILKHNFTSQSQK  102
JBBCAEG-19-0032_JBP_33490   60  KMGCPV----DL------LSIDHSRIPVAQGLI----EHLRGSASIATATQVFQAISCFFDWIDDQPRR-  114
WP_014386728.1_Vda04Ya311   50  --GAPNYSQQGKKIKGVDYIPVAGREAFVRDAYRLLRTDFN-----RTKQNHFEQLRLYIRWMDREHRK-  111
Consensus_aa:                   ...hss.....b......b....sRbshhp.h.pbhppchp.t.t.so....@.plp.@lp@hD.p..p.
Consensus_ss:                                           hhhhhhhhhhhhhhhhh   hhhhhhhhhhhhhhhhhhhhh


Conservation:                         9             7                                   7           7
WP_013855472.1_VagAT68554  111  -----PF------SQGGYLSYVGNSGQLWRLVNAANEPKRYHFQYHDGEEAGLLEDTAFIQKTLLDILLT  169
BCN27154.1_6283            107  -----PF------SESGYLKYAGNDGELRHRIKMFTPSKRL-WEYNHGDELSKKESSASTVLSTLRTALE  164
WP_017821119.1_ValAT17749   98  -----AF------TQSSLEGYM---VHLQTRVMLGE----------------LKSSTYKNRRANMVTLFS  137
BCN27178.1_6945            103  RKSKVEF------SPESCVNYI---QAVWLSQRTTGVG-------LHGK--PIKAKTLGWLASNYNSILK  154
JBBCAEG-19-0032_JBP_33490  115  -----CFFDDVAAMQKGYGDYT---QHLLHRMNSSGV---------NGK--PIKKTTASQYQAGARKVLM  165
WP_014386728.1_Vda04Ya311  112  PINGDYF------APDLYNAFM---DYHQDKCNRGEQ----------------SLSTWSNAKRIVSFFLK  156
Consensus_aa:                   ......F......s.pth.sYh.....lbpphp.s.................lp.sTh....s.hp.hLp
Consensus_ss:                                 hhhhhhhh   hhhhhhhhh                   hhhhhhhhhhhhhhhhh


Conservation:                                                        7
WP_013855472.1_VagAT68554  170  VL-DFD----VSEWQATLKPFSIKNSESSTQPY-TSSEWYALVRRTQLFFFSLATQLIAFKEENPEAPPP  233
BCN27154.1_6283            165  WC-GLP----VSDWARHHRGFA--RENEPFKGY-SDDEETLLVTRLDTLFFTLAPQLIAAKENNT--LLP  224
WP_017821119.1_ValAT17749  138  RYLDLP----HYYFNNVVIMDN--SDRESYESY-TQSDLKQLLPFLRRLFNQTYSQFIKNPE--------  192
BCN27178.1_6945            155  KF-GFG------DIPKATRNLDTASASLDSNNY-TKKELKRIARALLSDRKLLYKEYLDKT---------  207
JBBCAEG-19-0032_JBP_33490  166  LTTGLS-EPEVKGIATHISQKDGQSRHINLALP-NADEQARTFAVLVNFIDEAHRILVQGGA--------  225
WP_014386728.1_Vda04Ya311  157  AY-NRPIEAKQLILIK---GTK--KQANSHKGIDVVGELKPLVRRFIAAFAGFRRHFLEGT-------KP  213
Consensus_aa:                   .h.shs.......h.phh...s..ppp.shpsh.s.sEb..lh..h..hh..hhpphl............
Consensus_ss:                   hh         hhhhh                   hhhhhhhhhhhhhhhhhhhhhhhh


Conservation:
WP_013855472.1_VagAT68554  234  QCLEGI--------VVDRNNGRDITITLRGKD--------------------------------------  257
BCN27154.1_6283            225  ETLPLIISLGLHEETLHIPTSLETRFASKNGTA-------------------------------------  257
WP_017821119.1_ValAT17749  193  ---AHI----------QAHKSTPTMIFEWQGRE-------------------------------------  212
BCN27178.1_6945            208  ------------------------------ISE-------------------------------------  210
JBBCAEG-19-0032_JBP_33490  226  ---FPL--------CLASPNGESVYLYSEQVDTKKAKAANFSLAPFLANSPTFPTWGEIKAHFGLEGGSA  284
WP_014386728.1_Vda04Ya311  214  SLHPLWSEELFNQ-QAE-KN--GWSVKKRNQVKGTFKRTV------------------------------  249
Consensus_aa:                   .....h.............p..sh.l.p.p.sp.....................................
Consensus_ss:                        h        eee     eeeee


Conservation:                             6                           7       9  7               6
WP_013855472.1_VagAT68554  258  -----------GSVSPF---------------NQCMAAAYTLFAYYTAFNDTVIKDVRHPI-KVITSKVE  300
BCN27154.1_6283            258  ----------VNAGAAF---------------NLTMGAAYHLMCFFTSQNDSNIRNIAHPI-HVHTEDR-  300
WP_017821119.1_ValAT17749  213  ----------YKLCGGI---------------TKMMCAATYLLAYYTYANTTDLFRLKQPN-NASIS---  253
BCN27178.1_6945            211  ----------YKREIAF---------------NRLICNAVFLYVYYTAVGQAEALNTFVVEDSWSID---  252
JBBCAEG-19-0032_JBP_33490  285  ATQKQRATYDYRRRIYLSNNDNLRSCLRQRIGSHAVAAGMLAFIAATSCNFSVAKNLEVD--LLEIV---  349
WP_014386728.1_Vda04Ya311  250  ----------VGENAAR---------------NHFSRLAAMIAFCFTGQNTTPLLNLRFSD-IRFTA---  290
Consensus_aa:                  ...h.hs...
Consensus_ss:                              hhhhhh               hhhhhhhhhhhhhhh    hhhhhh      eeee


Conservation:                               9 9                                  6           76      7
WP_013855472.1_VagAT68554  301  GRTSKTAQVRAYKSRAS-KDVKALFAGSDENLHSEAIEKEAGFIVAEINKRDAVGKIDGITFIQTLELLS  369
BCN27154.1_6283            301  DKSLQVVKVSSYKPRSS-NEVDALLVGE----------------SFDVNKR------DGVKFIKLLERLS  347
WP_017821119.1_ValAT17749  254  -TGETWYTMPAFKRRAF-KTIQVEIGEH----------------ELEIPKY-------AMTFFDKLLNAS  298
BCN27178.1_6945            253  KAGGNRISVKGLKTRGY-KEESRTFA---------------------PRAV-------SKTFFEEHFELS  293
JBBCAEG-19-0032_JBP_33490  350  -PTSQGNRLSGTKGRARGKKVSPEFG-----------------------AR-------FAPVFKKYLDLR  388
WP_014386728.1_Vda04Ya311  291  -KSDGKVYFDMTKARANYLGFDTSLG---------------------FHKK-------TQAFFHQWLEVS  331
Consensus_aa:                   ..s.p.hphpthKsRt..pphps.ht......................p.........s.sFhcphbplS
Consensus_ss:                       eeeeeeee         eeee                   ee   hh       hhhhhhhhhhhh


Conservation:
WP_013855472.1_VagAT68554  370  KSYSND-PFDTLIYFLDKEGE---KDKVNVSHALLKLS--QNLNLLSSRRGELTEHLIKTYTDIVENQNM  433
BCN27154.1_6283            348  KLYGNSEDGSELLFTLNNNSE--VSNAFNLRELNKALV--NKLHLLSPHRAGNLPWFK-ELFYTYRNQQS  412
WP_017821119.1_ValAT17749  299  RIISTN-EDATLLQTIASRKVQ-KLQTSRLQSFLKKWV--EKHFSFTDQMGRR-----------------  347
BCN27178.1_6945            294  KLNAVDLGLDKHYLFTRRNGN--QPTPMNLNTYIASLM--TRSVLLQ-----------------------  336
JBBCAEG-19-0032_JBP_33490  389  KWVLNGAESTLVFPFFSPQYGF-VSLTGHQINRTKILF--NKALPHT-----------------------  432
WP_014386728.1_Vda04Ya311  332  MVLQKNAGTEWVFPYFRKSGEIQGCVDAGQTTPQKSINKCTKILGLA-----------------------  378
Consensus_aa:                   +hh.ss...s.lh.hhppp.......s.pbpp..b.l...p+hh.hs.......................
Consensus_ss:                   hhh       eeeeee        h hhhhhhhhhhhh  hhh


Conservation:
WP_013855472.1_VagAT68554  434  PIFNWSQREDGARTMNKQVIYLNKRTATKRSTPIAYAVFSCMADVSLRNALIPLHYAEKDANGEITVSFK  503
BCN27154.1_6283            413  ITLKKVTNHLGRTVVHKEVQEVSKTKAGQGATNSAYCILSCYTDLPYKGVLLPLTYSEIDSDGNVTVSFK  482
WP_017821119.1_ValAT17749       ----------------------------------------------------------------------
BCN27178.1_6945            337  -TMKA-----------------------------------------------------------------  340
JBBCAEG-19-0032_JBP_33490       ----------------------------------------------------------------------
WP_014386728.1_Vda04Ya311       ----------------------------------------------------------------------
Consensus_aa:                   ......................................................................
Consensus_ss:                    eee


Conservation:
WP_013855472.1_VagAT68554  504  YVDGSEGEFTVTAKHLPFLQLVERHA---ATRNPLPKERSLGGIGRRSNATKPPFLLPLGTKSMTYQWQE  570
BCN27154.1_6283            483  YRSGETGYFKAPACDLTLIKDIERYAKEKANKQAKKYKRLLLTRSS---------------NNIPYDW-E  536
WP_017821119.1_ValAT17749       ----------------------------------------------------------------------
BCN27178.1_6945            341  ----ANPDF-------------------------------------------------------------  345
JBBCAEG-19-0032_JBP_33490       ----------------------------------------------------------------------
WP_014386728.1_Vda04Ya311       ----------------------------------------------------------------------
Consensus_aa:                   ......................................................................
Consensus_ss:


Conservation:                                                7            6          7          9
WP_013855472.1_VagAT68554  571  GEVPISASMLSYCGIGYGDYFLNINSRRIRVTHSDLEYKPE-ERGMTAQKILQHSIDTAD--KKYRNGHP  637
BCN27154.1_6283            537  GISPISANLMKRWSVEPNHYYLSLQSSRWREMSSNQAYAEG-G-VQAVQSLLQNKRDTIE--RSYINGLP  602
WP_017821119.1_ValAT17749  348  ---------------------LRPSISRFRETGCQLTSYHQ-G-EMVNNLMLNNTPNTRK--RHYSEGNK  392
BCN27178.1_6945            346  ----------------------SLTCERLKSSIKQYAEEKL-G-RQGAMESNRNTSESTWNNSDYSKNSK  391
JBBCAEG-19-0032_JBP_33490  433  ---------------------SWVTPTQWRKGVSYQYVKLSGGDLALTAEKLGNTEATLR--QSYSRPAL  479
WP_014386728.1_Vda04Ya311  379  ----------------------HVTPSKLRQTKIDTLMKVT-EDIWLVSMSANNSIEVVA--INYSDGNE  423
Consensus_aa:                   ......................plssp+hRps..pbh.b......bhh.b.hpNo.poh...ppYpps..
Consensus_ss:                                            hhhhhhhhhhhhhhhh   hhhhhhhh   hhhhh  hhh    h


Conservation:                                              6                                  9   9
WP_013855472.1_VagAT68554  638  VQNNKQVSQGLMALSRIAEG--ETRNEAIESVKAELKIPVLEYEVWKKRNQ------PTNPNG-ISCDG-  697
BCN27154.1_6283            603  SLNKVILSQGIEVIENLFDN---DLEKAKDAVAKRRGIPMLTYEESEKKRK-------TNPNG-IVCDG-  660
WP_017821119.1_ValAT17749  393  LANNGMMQDTMSIREEQVKS-----RVNTKQAQKNLSIEVLVIEEENKINL---PNLSRTSNG-GSCAA-  452
BCN27178.1_6945            392  ALAHRELAFGTATLFALGSNPTGGVTAAIATTKAQHC-EVLSSEEVDALRASSSEPVDAIANG-GVCKG-  458
JBBCAEG-19-0032_JBP_33490  480  EDFAAEMTVFFELMHKAA-------------IDRTRSVEHIPVRIVDEIRL-------DAVTGIGSCEKA  529
WP_014386728.1_Vda04Ya311  424  SDHRTSLAANNEALYDFSKNG-TDPHEAANKSKFNHANVLNGYDYKRLRKE-ETENDTQTPLG-VRCKD-  489
Consensus_aa:                   ...p.bhs.sh.hh.phsps.....p.Ah.php.p.t..hlsh-b.cb.pb.......psssG..sCcs.
Consensus_ss:                   hhhhhhhhhhhhhhhhhh      hhhhhhhhhhhh      hhhhhhh


Conservation:                                    9     7           9     9  9          69  779      9
WP_013855472.1_VagAT68554  698  --KIDLNSEKDWHYAARKFAEEKGIIKEGD--DITCYQYDLCPFCKSAKLVDDPYAIYKLLSFLDVFSEA  763
BCN27154.1_6283            661  --KQAM--------IDGKNTQRETNYALDA--DLPCAEFDMCHKCQSAKALDEVDAIYKLISYIDVLKEG  718
WP_017821119.1_ValAT17749  453  --PFGEKSQ-----KYTKKAQTQGLAKEGE--RLACADLLGCFGCPDQVIVQSVSDIWCLLSFKSCVEES  513
BCN27178.1_6945            459  --EDVPQKR-----EFQKSNLETGLLDDDDVKNTACGYVIKCFMCRNFAVVDEVHDIWRLLSFEFRLNEA  521
JBBCAEG-19-0032_JBP_33490  530  AETAPRR--------------AQGFTAQAP--APACGDPETCLFCEFYAVHADEQDIRRLLSLRYLIE-A  582
WP_014386728.1_Vda04Ya311  490  --SSQGVAV-----RIKKNLENMGVKQPEK--EQRCTDFLGCFECHYHRLVSEVEDIWLMLSFNDTLQEM  550
Consensus_aa:                   ..p.s..........hpKp.bppGhh..s...p.sCsph..ChbCp.h.lls-spsI@.LLS@..hlpEt
Consensus_ss:                        hhhh     hhhhhhhh             hhhhhh      eeee  hhhhhhhhhhhhhhhhh


Conservation:                                       9                    7       6  7
WP_013855472.1_VagAT68554  764  IDRY------PERASVIQSKIERFQEHLDDLP---FETIEKAEDLLEEKGRYPLFHSLS-SVTQFL----  819
BCN27154.1_6283            719  LDMF------PDSKGDALEKIKAFECTLDEAS---NDVFDEAMKKFNTQGRHPRVSIDH-AILSM-----  773
WP_017821119.1_ValAT17749  514  LFLHLDVSHYKQNFENIILFINQ--KILPNIN---KQILRKAETKLDDDGLHPAWDDSE-SVLSLIPKSP  577
BCN27178.1_6945            522  VAAHKSLDHFIKNFSEVKSAIRDIKKRFK------KRNLKAAEKYLERQGCHPLWDED--SIQDIFKG--  581
JBBCAEG-19-0032_JBP_33490  583  IKHKQPIDHWQTKFSPTLHRIDEVLSAIQDADGRIKPTVTRVRDEVASGAFDAFWSIHFDTLVTVGAVT-  651
WP_014386728.1_Vda04Ya311  551  KAYPAINSLPTDKFYKLCNTIESILKDFKEVA---PDNYTRAQEMH-SEAPHPLYSDGY-SLMDLLETF-  614
Consensus_aa:                   l.h........pph..hbp.Icphbp.hpphs....pshpcApcbh.ppt.cPh@s....olhsh.....
Consensus_ss:                   hh      hhhhhhhhhhhhhhhhhhhhh      hhhhhhhhhhhhh         hh hhhhh


Conservation:
WP_013855472.1_VagAT68554       ----
BCN27154.1_6283                 ----
WP_017821119.1_ValAT17749  578  AEVR  581
BCN27178.1_6945                 ----
JBBCAEG-19-0032_JBP_33490       ----
WP_014386728.1_Vda04Ya311       ----
Consensus_aa:                   ....
Consensus_ss:
```

**FIG S2.** Secondary structure alignment of JBP_33490 product of JBBCAEG-19-0032 with known Tfp homologs. The alignment was generated using PROMAL3D. e: sheet; h, helix.
