## Supplementary material for "Nationwide genome surveillance of carbapenem-resistant *Pseudomonas aeruginosa* in Japan": Fig. S3

```
Conservation:                   9                    7  97 77                        6    7  797   7
WP_013855470.1_VagAT68554    1  MTDRTE--------LSSTEKQLREALKRLIDKQPTHR---ELKRKLSANKL-KIVVANVEKEAGLSNGAA   58
WP_014386726.1_Vda04Ya311    1  MNQETLRALVQEEKTLAAEEKLHKALARLQNGKPLRT----------KAKG-RLTLNKINNEAGLGRSYI   59
BCN27180.1_6945              1  MA--------------TVATKLENAFKALLKEGK------------------KISPYAVERRAGVSNGSL   38
WP_017821121.1_ValAT17749    1  MSK-----------SENTLLKLEAALERIKQGKPKRI----------PTHR-KLSVRAVEEEANLGNGSG   48
BCN27152.1_6283              1  MS-------------QTTKKKLIDALERLLSGDVAKLTSRELRNKA-RKGKLKINNSNVEKEAGLSVGAL   56
JBBCAEG-19-0032_ JBP_33510   1  MS---------------ALENYIAALQRLIAGKPQNV----------PKGS-AINKDTVALEAGRKRGSI   44
Consensus_aa:                   Ms..............shbppL..ALpRLbp.ps.p................+ls..sVppEAGltpGth
Consensus_ss:                                  hhhhhhhhhhhhhhh                      eeehhhhhhhh     hh


Conservation:                                6                                                776
WP_013855470.1_VagAT68554   59  KRY----PETKKMIEGAEAERIHGSSDVSGEVVRAHPIYIKARE----DLEK---AREEVKKLKAELEQK  117
WP_014386726.1_Vda04Ya311   60  HKF----KEFVDNVANPAIEAYNETLDTP----KPSEIDKAS-TEDMSEIDRLKDELRKQIELKEAY---  117
BCN27180.1_6945             39  KNH----IVLLEKVLAEKEKYATDPTNVG--VVKARANKAKAKV----SKDKYQEVLDKNEKLRAEN---   95
WP_017821121.1_ValAT17749   49  YYY----PDFVEKVKQTKGEIATGKGGIV----QPEILTVRTKLK---EQKR------IKDNYKAKYE--   99
BCN27152.1_6283             57  RRH----NDVILMIKNKSLEVQVAQDETS----DSPIEILLKEIK---SIRS---EKIREKNLKEEYY--  110
JBBCAEG-19-0032_ JBP_33510  45  KKSRAGNAELIAAIEAAAAAQQEKSGPTA-------AQDATKQKA---LKRA---AQAQLGSLKEDY---   98
Consensus_aa:                   +p@.....-hl..lbs...c..p.psshs....csp.b.h..p.....pbcp....b.p..pLK.ch...
Consensus_ss:                   hhh    hhhhhhhhhhhhhhhh          hhhhhhhhhhhh   hhhh   hhhhhhhhhhhhh


Conservation:                                 66                            6
WP_013855470.1_VagAT68554  118  DGRLAHYKELLKSQAARMHQMNVAMWNDIPEEKKHVEVIIDVQDMG---------SQD---NILAFKKRE  175
WP_014386726.1_Vda04Ya311  118  -------RTERDEALATNDELEVLN----------KSLMFRVYELQ-------QELTD---SVAEYEGYK  160
BCN27180.1_6945             96  ------------------EQFKGEL----------KVMADKVAQMTWELHRYKTKTRKDSTNVHNLKA--  135
WP_017821121.1_ValAT17749  100  -----AEREKLALFAAAQHHLNDKL----------VQALARIDDLEY--ENA--ELRE---ELAKLKRSK  147
BCN27152.1_6283            111  -----GEAQNHQDALAAQAATHVKV----------VQE---LMDMLHESERE--KAMD---KVVNARPDN  157
JBBCAEG-19-0032_ JBP_33510  99  -----ELALTRIASLVHENHTLKMQ----------IKE---LVE-----EKE--WSKH---KVVKMNEAK  140
Consensus_aa:                   ..............hh..pphpsb............p....l.-h.........c.pc...plhphc..c
Consensus_ss:                        hhhhhhhhhhhhhhhhhhhh          hhhhhhhhhhhh  hhh  hhhh   hhhhhh


Conservation:
WP_013855470.1_VagAT68554  176  KLN-----  178
WP_014386726.1_Vda04Ya311  161  K-------  161
BCN27180.1_6945                 --------
WP_017821121.1_ValAT17749  148  VVPIK---  152
BCN27152.1_6283            158  IVEGHFRK  165
JBBCAEG-19-0032_ JBP_33510 141  RP------  142
Consensus_aa:                   ........
Consensus_ss:
```

**FIG S3.** Secondary structure alignment of JBP_33510 of JBBCAEG-19-0032 with known Srap homologs. The alignment was generated using PROMAL3D. e: sheet; h, helix.
